# Retinal metabolism: Evidence for uncoupling of glycolysis and oxidative phosphorylation via Cori-, Cahill-, and mini-Krebs-cycle

**DOI:** 10.1101/2022.06.20.496788

**Authors:** Yiyi Chen, Laimdota Zizmare, Victor Calbiague, Lan Wang, Shirley Yu, Friedrich W. Herberg, Oliver Schmachtenberg, François Paquet-Durand, Christoph Trautwein

## Abstract

The retina consumes massive amounts of energy, yet its metabolism and substrate exploitation remain poorly understood. Here, we used a murine explant model to manipulate retinal energy metabolism under entirely controlled conditions and utilized ^1^H-NMR spectroscopy-based metabolomics, in situ enzyme detection, and cell viability readouts to uncover the pathways of retinal energy production. Our experimental manipulations resulted in varying degrees of photoreceptor degeneration, while the inner retina and retinal pigment epithelium were essentially unaffected. This selective vulnerability of photoreceptors suggested very specific adaptations in their energy metabolism. Rod photoreceptors were found to rely strongly on oxidative phosphorylation, but only mildly on glycolysis. Conversely, cone photoreceptors were dependent on glycolysis but insensitive to electron transport chain decoupling. Importantly, photoreceptors appeared to uncouple glycolytic and Krebs-cycle metabolism via three different pathways: 1) the mini-Krebs-cycle, fueled by glutamine and branched-chain amino acids, generating N-acetylaspartate; 2) the alanine-generating Cahill-cycle; 3) the lactate-releasing Cori-cycle. Moreover, the metabolomic data indicated a shuttling of taurine and hypotaurine between the retinal pigment epithelium and photoreceptors, likely resulting in an additional net transfer of reducing power to photoreceptors. These findings expand our understanding of retinal physiology and pathology and shed new light on neuronal energy homeostasis and the pathogenesis of neurodegenerative diseases.

**Graphical abstract.**
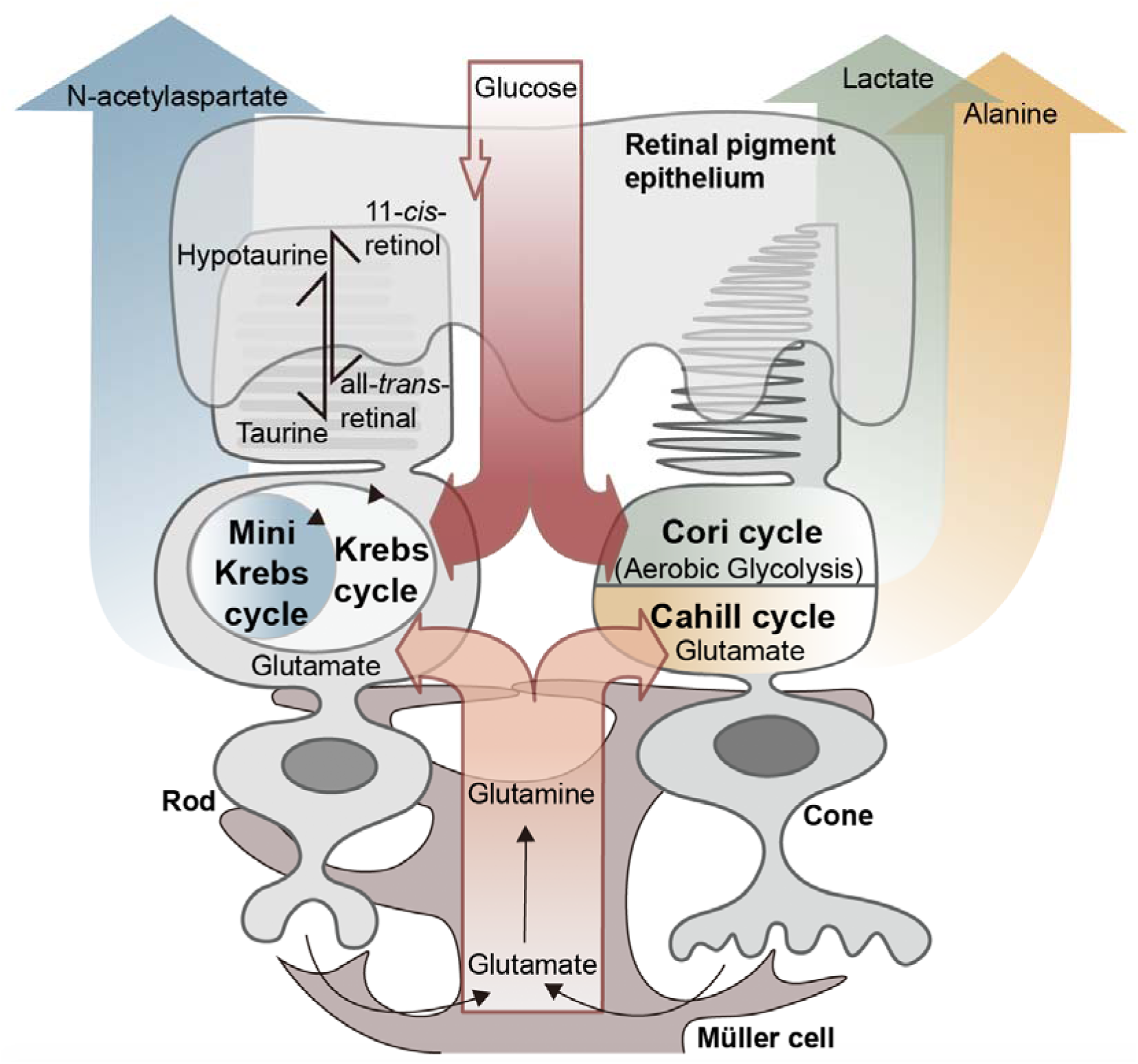
Retinal photoreceptors employ both glucose and glutamate as fuels. While rod photoreceptors rely strongly on oxidative phosphorylation and the N-acetylaspartate producing mini-Krebs-cycle, cone photoreceptors rely on the lactate-producing Cori cycle and the oxidative, alanine producing Cahill cycle.

**Highlights:**

- The retina utilizes a complex energy switchboard consisting of the Krebs cycle, mini-Krebs cycle, Cahill cycle, and Cori cycle.
- Mini-Krebs cycle runs more efficiently than ‘full’ Krebs cycle.
- Alanine transaminase decouples glycolysis from the Krebs cycle.
- Lactate, alanine, and N-acetylaspartate are distinctive energetic pathway signatures.

## Introduction

The retina is the neuronal tissue with the highest energy demand (1, 2). Paradoxically, the mammalian retina is thought to strongly rely on energy-inefficient glycolysis, even though oxygen is available and high-yield oxidative phosphorylation (OXPHOS) possible (3). This aerobic glycolysis releases large amounts of lactate, as already reported in the 1920s by Otto Warburg (4). High retinal energy demand is linked to the extraordinary single-photon sensitivity of photoreceptors (2, 5, 6), and to lipid synthesis for the constant renewal of photoreceptor outer segments (7, 8).

The retina harbors two types of photoreceptors: rods, which exhibit remarkable light sensitivity and enable night vision; and cones, which work in daylight and allow color vision. Cones spend around twice as much energy as rods (9). Photoreceptors are not connected to the vasculature and are nourished by other retinal cell types, including retinal pigment epithelial (RPE) (10) cells or Müller glial cells (MGC). Moreover, photoreceptors can experience changes in energy demand on a millisecond timescale, from very high in the dark, to 4-10 times lower in light (5).

Glycolysis provides for rapid but inefficient adenosine triphosphate (ATP) production, while Krebs-cycle and OXPHOS are very efficient but much slower. Since glycolysis and Krebs-cycle are metabolically coupled through pyruvate, it is unclear how photoreceptors adapt to sudden and large changes in energy demand, and what energy substrates they may use. A recent hypothesis proposed that photoreceptors use predominantly aerobic glycolysis, with the resultant lactate utilized by the RPE and MGCs for OXPHOS (11). Such an arrangement would be reminiscent of the Cori-cycle, in which glucose and lactate are cycled between skeletal muscle and liver (12). However, this concept contrasts with the high density of mitochondria in photoreceptor inner segments, which strongly suggest an extensive use of OXPHOS.

Alternative pathways for ATP-generation include the Cahill-cycle, where pyruvate is transaminated using, for instance, glutamate to yield alanine and α-ketoglutarate (13, 14). The advantage of Cahill-over Cori-cycle is that reducing equivalents (i.e. NADH) gained during glycolysis are preserved and that α-ketoglutarate can yield additional ATP in mitochondrial OXPHOS. A second alternative may be the so-called mini-Krebs-cycle, in which a transamination of oxaloacetate with glutamate produces aspartate and α-ketoglutarate (15). This shortens the “full” 10-step Krebs-cycle to a significantly faster 6-step cycle. A point that Cori-, Cahill-, and mini-Krebs-cycle have in common is that they do not require the pyruvate-coupling between cytoplasmic glycolysis and mitochondrial OXPHOS, allowing both processes to run independent of each other. Whatever the case in the retina, the question as to which pathway is used has obvious ramifications for disease pathogenesis, including for diabetic retinopathy, age-related macular degeneration, or inherited retinal diseases, such as retinitis pigmentosa.

Here, we studied the expression of energy metabolism-related enzymes in different retinal cell types, using organotypic retinal explants maintained in serum-free, fully defined medium. Metabolic functions of the RPE were investigated by culturing retina with and without RPE. As readouts, we correlated enzyme expression with quantitative proton nuclear magnetic resonance (^1^H-NMR) spectroscopy-based metabolomics and cell death. Notably, interventions in energy metabolism caused selective photoreceptor cell death, while leaving other retinal cell types essentially unaffected. Metabolomic analysis and localization of key enzymes identified important pathways and shuttles altered, explaining the strong interdependence of the various retinal cell types, and opening new perspectives for the treatment of neurodegenerative diseases in the retina and beyond.

## Results

### Retinal expression patterns of key energy metabolism related enzymes

We used immunofluorescence to assess expression and cellular localization of enzymes important for energy metabolism in wild-type mouse retina in vivo (Figure 1). The outermost retinal layer is formed by RPE cells expressing RPE-specific 65kDa protein (RPE65), dedicated to the recycling of photopigment, retinal (16). Hence, RPE65 immunolabeling revealed the RPE cell monolayer (Figure 1A). Immunofluorescence for glucose-transporter-1 (GLUT1) showed strong labelling on basal and apical sides of RPE cells (Figure 1A), in line with previous literature (3, 17). GLUT1 expression was not detected in rods or cones (Supplementary Figure 1A, B). Instead, glucose uptake in the outer retina seems to be mediated by high affinity/high capacity GLUT3 (18), strongly expressed on photoreceptor inner segments (Supplementary Figure 2A). Pyruvate kinase is essential to glycolysis, catalyzing the conversion of phosphoenolpyruvate to pyruvate. While pyruvate kinase M1 (PKM1) was expressed in the inner retina (Supplementary Figure 2A), PKM2 was found in photoreceptor inner segments and synapses (Figure 1A). Expression of mitochondrial cytochrome oxidase (COX) largely overlapped with PKM2 (Figure 1A).

**Figure 1.**
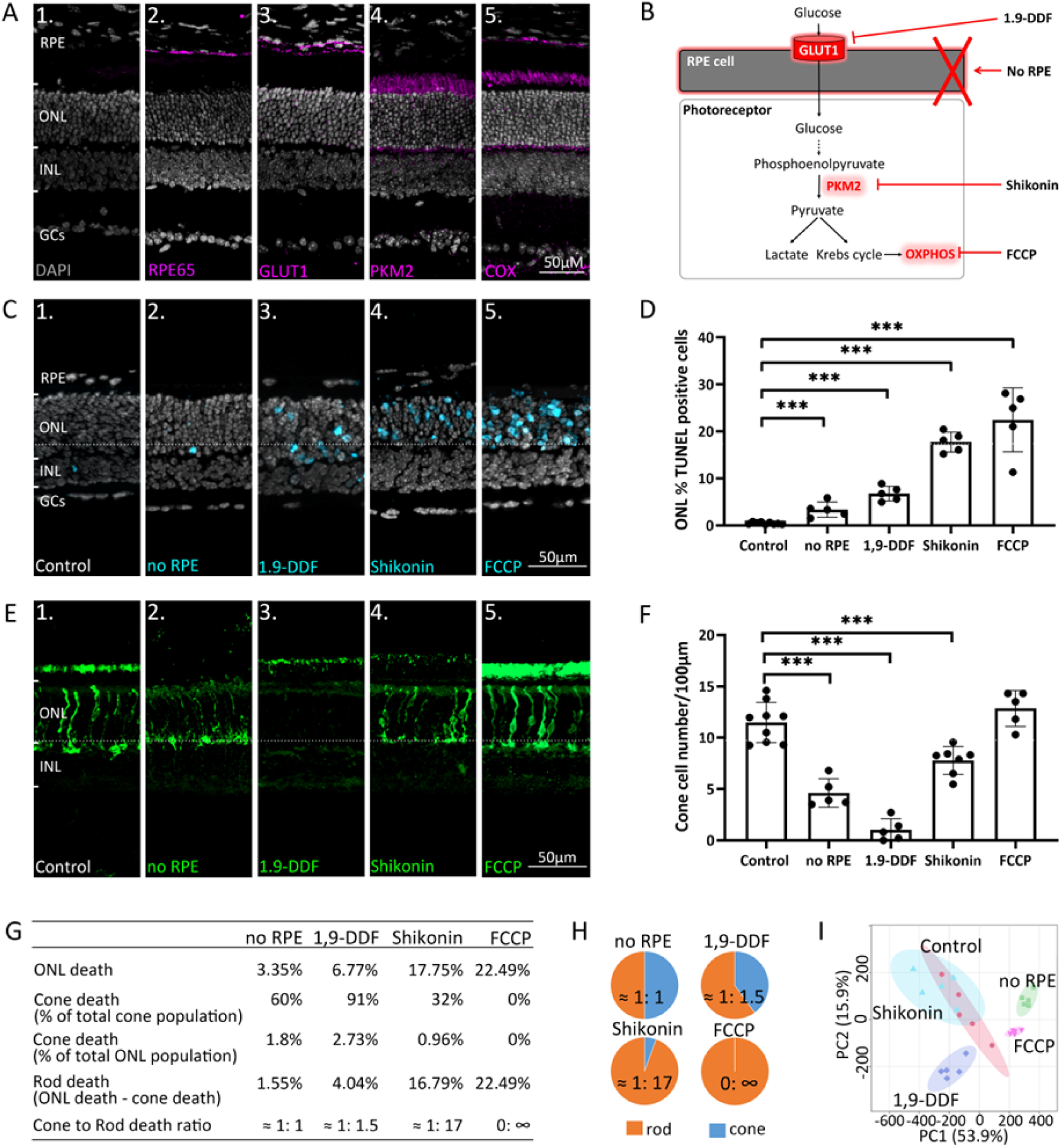
Manipulating energy metabolism differentially affects rod and cone photoreceptors. (A) Immunofluorescence staining of *in vivo* retina (magenta): (A1) negative control, (A2) RPE65, (A3) glucose transporter-1 (GLUT1), (A4) pyruvate kinase M2 (PKM2), (A5) mitochondrial cytochrome oxidase (COX). (B) Overview of experimental manipulations. Organotypic retinal explants were cultured for 6 days in vitro with or without RPE or treated with 1,9-DDF, Shikonin, or FCCP. (C) TUNEL assay performed on in vitro retina (cyan) marking dying cells in the five experimental conditions. (D) Quantification of TUNEL-positive cells in outer nuclear layer (ONL). Data represented as mean ± SD. (E) Cone-arrestin labelling (green) in the ONL. (F) Quantification of arrestin-positive cones. Data represented as mean ± SD. (G) Table giving percentages of cones and rods lost for each treatment. (H) Pie charts illustrate cone to rod cell death ratios. (I) Principal component analysis (PCA) of retinal samples investigated with ^1^H-NMR spectroscopy-based metabolomics. Dots in graphs represent individual retinal explants. Statistical testing: One-way ANOVA with Tukey’s post-hoc test. Asterisks indicate significance levels; p-values: *** < 0.001, ** < 0.01 * < 0.05. RPE = retinal pigment epithelium; INL = inner nuclear layer; GCs = ganglion cells.

### Challenging energy metabolism reduces rod and cone photoreceptor viability

To dissect retinal energy metabolism, we selectively manipulated key pathways (Figure 1B) using organotypic retinal explants (19) (Supplementary Figure 2B). Retinal cultures were prepared with the RPE cell layer attached to the neuroretina (control) or without RPE (*no* RPE). The *in vivo* expression patterns of enzymes important for energy metabolism (Figure 1A) were comparable in the *in vitro* situation (Supplementary Figure 2C). In control, the rate of cell death in the outer nuclear layer (ONL), as assessed by the TUNEL assay (Figure 1C, D), was low (0.46%±0.22, n=9). The *no* RPE condition displayed significantly increased photoreceptor cell death (3.35%±1.62, n=5, p<0.001). Blocking RPE glucose transport with the selective GLUT1 inhibitor 1,9-dideoxyforskolin (1,9-DDF) (18) further increased ONL cell death (6.77%±1.57, n=5, p<0.0001). The inhibition of glycolytic PKM2 with Shikonin (20), and OXPHOS disruption with the electron chain uncoupler carbonyl cyanide-p-trifluoromethoxyphenylhydrazone (FCCP) (16) both caused a significant increase in photoreceptor cell death (Shikonin: 17.75%±2.13, n=5, p<0.0001; FCCP: 22.49 % ± 6.79, n=5, p<0.0001). All four experimental interventions mostly affected photoreceptors, not significantly reducing the viability of other retinal cell types (Figure 1C).

Remarkably, a differential effect on cone photoreceptor survival (Figure 1E, F) was observed as assessed by immunodetection of the cone-specific marker arrestin-3 (Arr3). Control retina harbored 11.47 (±1.95, n=9) cones per 100 µm of retinal circumference. *no* RPE retina displayed a significant decrease in cone survival (4.62±1.39, n=5, p<0.0001), while blocking RPE glucose transport with 1,9-DDF led to an even more dramatic reduction (1.02±1.09, n=5, p<0.0001). By comparison, inhibition of glycolysis with Shikonin had a relatively mild effect on cone survival (7.78±1.36, n=5, p<0.0001). Surprisingly, FCCP treatment did not cause cone death (12.84±1.75, n=5, p=0.22), compared to control.

We further calculated the percentages of dying cones and rods, assuming that 3% of all photoreceptors were cones (21, 22) (Figure 1G, H). In the four treatment situations the ratios of cone to rod death were: *no* RPE = 1:1; 1,9-DDF = 1:1.5; Shikonin = 1:17; and FCCP = 0: ∞. Cones were almost entirely depleted by the 1,9-DDF treatment but remained virtually unaffected by the FCCP treatment. Rods, however, were strongly and highly selectively affected by Shikonin and FCCP, while the *no* RPE condition had a relatively minor effect. These results highlight important differences between rod and cone energy metabolism.

### Experimental retinal interventions produce characteristic metabolomic patterns

We employed high-field (600 MHz) ^1^H-NMR spectroscopy-based metabolomics to study the metabolic signatures in the five experimental groups (Supplementary Table 1, Supplementary Figure 3). A principal component analysis (PCA) showed clear group separation (Figure 1I). Unbiased clustering of metabolite profiles revealed specific groups that were differentiated among the five experimental situations (Figure 2A). The greatest cluster overlap was seen in control and Shikonin groups. Significant clustering was observed in several amino acid sub-classes and energy metabolites. We found the strongest changes in the 1,9-DDF condition, where glutamate, aspartate, alanine, O-phosphocholine, and nicotinamide adenine dinucleotide (NAD) concentrations were particularly upregulated compared to control. Further, branched-chain amino acids (BCAAs), lactate, and threonine were upregulated in the FCCP and *no* RPE condition. Glutathione (GSH) and glutathione disulfide (GSSG) were relatively reduced by Shikonin treatment, with increased creatine and formate. Control and 1,9-DDF treatment displayed high levels of guanosine triphosphate (GTP), adenosine triphosphate (ATP), and π-methylhistidine, which were low in the other groups (Figure 2A).

**Figure 2.**
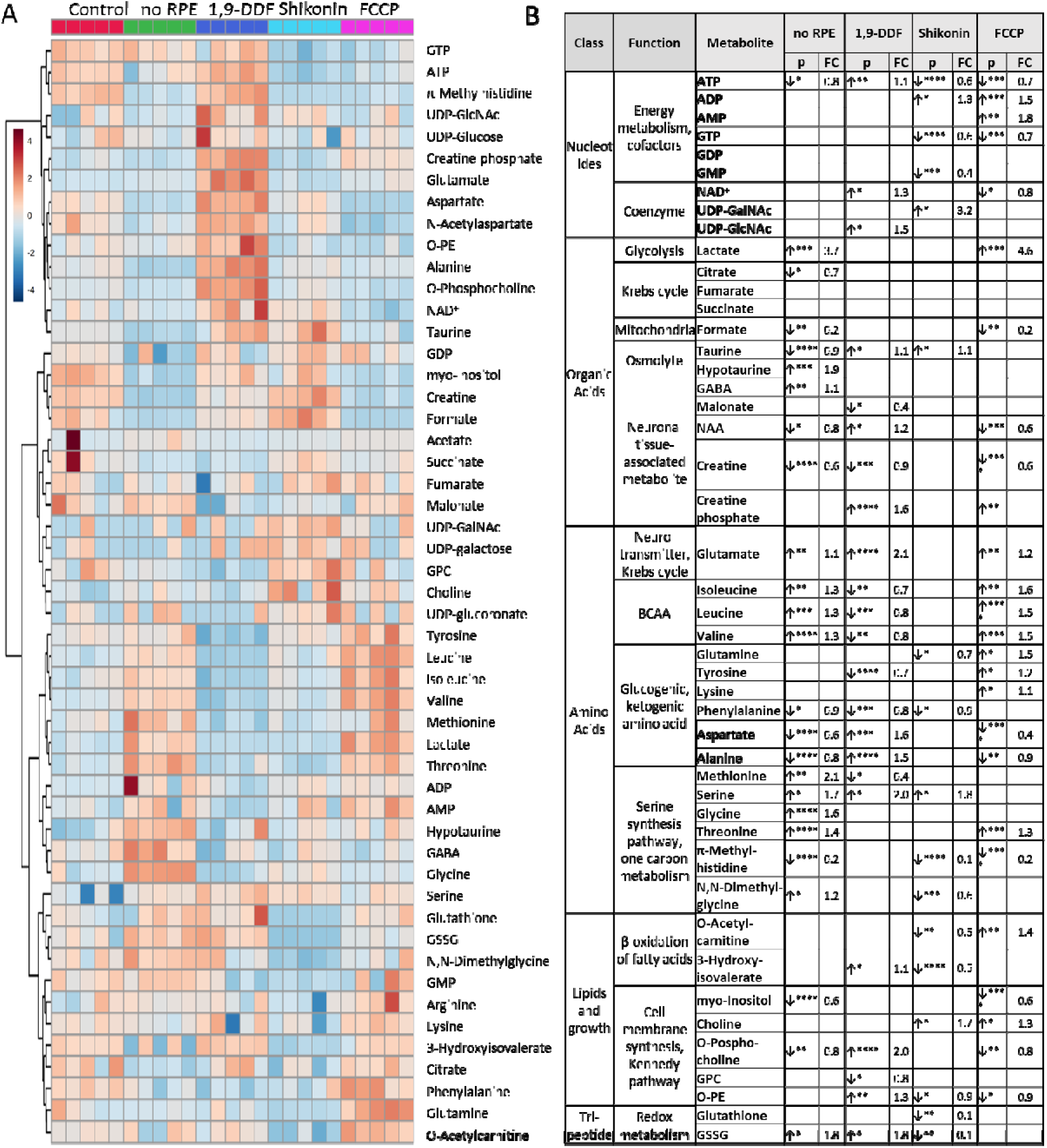
^1^H-NMR spectroscopy-based metabolomic analysis of retina subjected to interventions in energy metabolism. (A) Heatmap based on unsupervised hierarchical cluster analysis by Ward’s linkage, showing auto-scaled metabolite concentrations (red – high, blue – low) in five different experimental conditions: control - red, *no* RPE - green, 1,9-DDF - dark blue, Shikonin - light blue, FCCP purple (n = 5 samples per condition). (B) Metabolic profiles of each intervention were compared to control. Metabolites significantly changed in at least one experimental condition were grouped according to functions and pathways. Data show p-values and fold change (FC) over control. Statistical comparison: Student’s unpaired t-test (group variance equal); p-values: **** < 0.0001, *** < 0.001, ** < 0.01 * < 0.05. See also Supplementary Figure 3.

We then compared metabolite patterns for each treatment to control (Figure 2B). Metabolites were grouped according to class and function, including fundamental metabolic pathways, such as energy and cofactor metabolism, coenzymes, Krebs-cycle, neuronal tissue-associated metabolites and neurotransmitters, BCAAs, glucogenic and ketogenic amino acids (AAs), serine synthesis pathway, cell membrane synthesis, and individual features from redox metabolism.

To get an overall impression of treatment effects on energy metabolism, we assessed the adenylate energy charge of the retina under the various experimental conditions and the ratio of GSH to GSSG to investigate oxidative/reductive stress. The energy charge appeared to be reduced in the shikonin and FCCP condition, while the GSH/GSSG ratio of the tissue was not significantly changed from control in any condition (Supplementary Figure 4A, B). We then analyzed the spent culture medium at P15 to assess retinal glucose consumption and lactate production after two days of culture. As could be expected, 1,9-DDF treatment reduced glucose consumption compared to control, while FCCP treatment increased glucose expenditure (Supplementary Figure 4C). Conversely, lactate production was decreased in the 1,9-DDF group. However, in the *no* RPE condition and after FCCP treatment lactate concentrations in the spent medium were strongly and significantly elevated (Supplementary Figure 4D).

Afterwards, the metabolomic differences identified within the retina were explored in detail and related to the retinal expression of corresponding enzymes.

### Retina cultured without RPE displays strongly increased glycolytic activity

The RPE is tightly bound to the choroid and serves as interface between vasculature and neuroretina. Because of the strong adherence of RPE to the choroid, most prior studies on explanted retina were performed without RPE (4, 11). To understand the metabolic communication between neuroretina and RPE, we prepared organotypic retinal explants both with RPE (control) or without RPE (*no* RPE). Since most previous studies on retinal energy metabolism had used retina without RPE (4, 23), this comparison also allowed us to relate our data to those earlier results.

The metabolite profile of control was markedly different from retina cultured without RPE, with 25 metabolites exhibiting significant changes (Figure 3A, Supplementary Figure 5). Decreased ATP levels in the *no* RPE group, corresponded to increased lactate. The BCAAs isoleucine, leucine, and valine, and some of the glucogenic/ketogenic AAs threonine, methionine, glycine, displayed higher levels in the *no* RPE group compared to control. Alanine, aspartate, and taurine showed the opposite trend. Hypotaurine, 4-aminobutyrate (GABA), and glutamate were accumulated in retinal tissue, while membrane synthesis-associated metabolites, such as myo-inositol and O-phosphocholine, as well as creatine and N-acetylaspartate (NAA) were reduced in neuroretina. Moreover, higher levels of GSSG appeared in the absence of RPE, pointing towards an altered redox metabolism.

**Figure 3.**
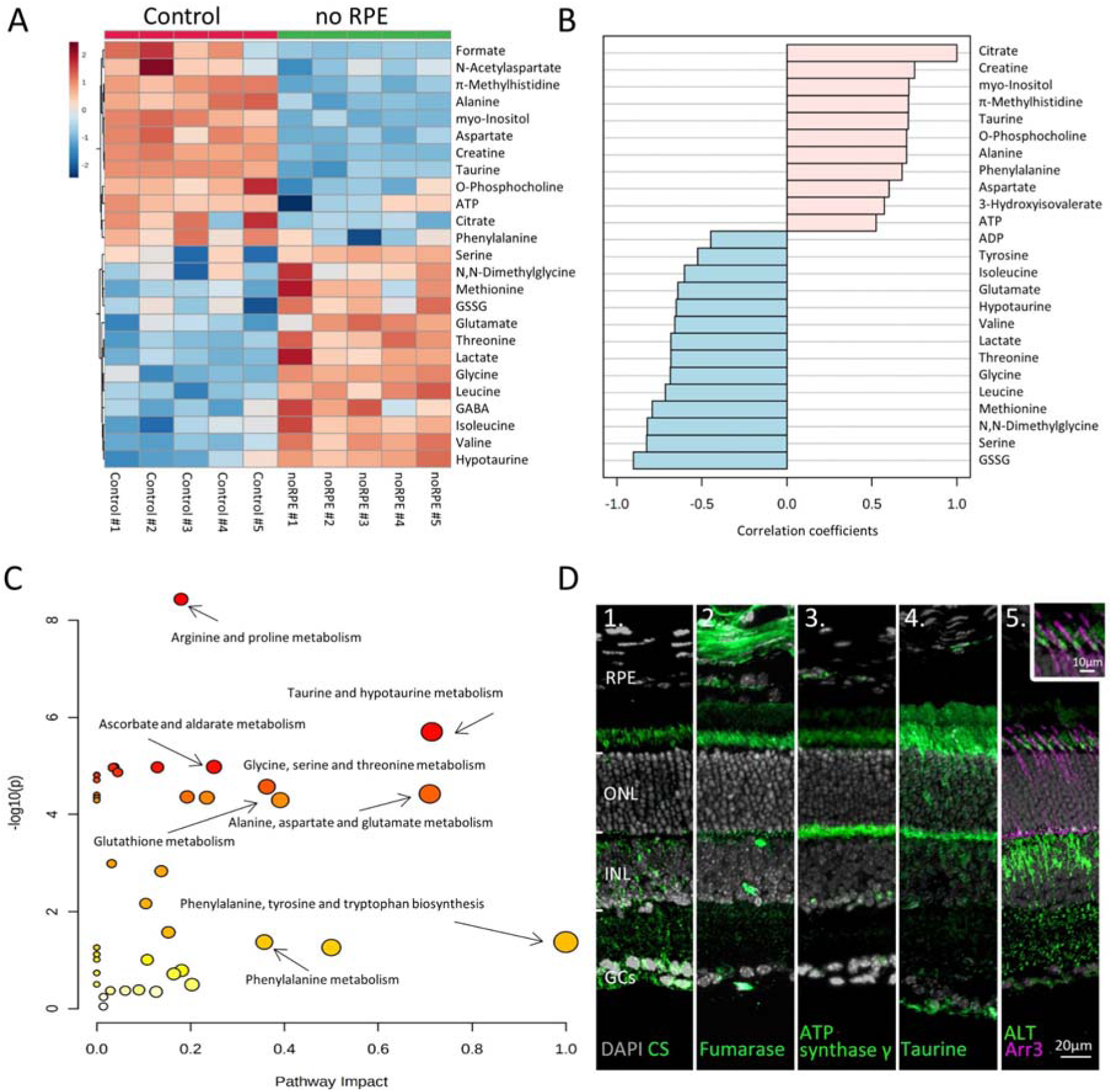
Absence of RPE dramatically changes retinal metabolism. (A) Heatmap, based on unsupervised hierarchical cluster analysis by Ward’s linkage, illustrating statistically significant changes for 25 metabolites (unpaired *t*-test, fold change (FC) > 1.2, raw p value < 0.05). (B) Pattern hunter for citrate, showing the top 25 correlating metabolites, correlation coefficient given as Pearson r distance. (C) Metabolic pathways most affected in *no* RPE condition compared to control, based on KEGG-pathway database. (D) Immunodetection of enzymes and metabolites (green) on *in vivo* retina. (D1) citrate synthase (CS), (D2) fumarase, (D3), ATP synthase γ, (D4) taurine, (D5) alanine transaminase (ALT). Co-staining for ALT and cone arrestin (Arr3; magenta) showed high expression in cone inner segments. DAPI (grey) was used as a nuclear counterstain. See also Supplementary Figure 5.

Pattern hunter (Pearson R) correlation analysis found citrate, a central Krebs-cycle metabolite, positively correlated with taurine, alanine, aspartate, and ATP (Figure 3B). Negative correlations with citrate included ADP, BCAAs, and lactate. A subsequent KEGG-based pathway analysis revealed changes in arginine and proline metabolism, taurine and hypotaurine metabolism, and alanine, aspartate, and glutamate metabolism in the *no* RPE situation (Figure 3C).

Immunofluorescence was used to localize enzymes likely responsible for the observed metabolic changes (Figure 3D). Citrate synthase (CS) and fumarase, key enzymes of the Krebs-cycle, were found mostly in photoreceptor inner segments and synaptic terminals. The γ-subunit of ATP synthase, essential for OXPHOS, was also localized to photoreceptor inner segments and synapses. Thus, low ATP levels in the *n*o RPE group likely resulted from decreased photoreceptor Krebs-cycle and OXPHOS activity. High levels of taurine were detected in photoreceptor inner segments and synapses, as well as in the ganglion cell layer (Figure 3D). Finally, the enzyme alanine transaminase (ALT), previously thought to be expressed only in muscle and liver, was found to be expressed in the inner nuclear and plexiform layers (Figure 3D). ALT was also strongly expressed in cone inner segments, as evidenced by co-labelling with cone-Arr3. This expression pattern and high alanine levels in the control strongly suggested Cahill-cycle activity in the retina (14). When ALT activity was inhibited with β-chloro-alanine, photoreceptor cell death was increased and notably cone survival was reduced, functionally validating the expression of ALT in the outer retina (Supplementary Figure 6A, B).

The patterns observed in the *no* RPE to control comparison indicated that retinal metabolism strongly depended on RPE-to-neuroretina interactions, including shuttling of metabolites such as hypotaurine/taurine. Notably, the accumulation of BCAA, glutamate, and lactate, concomitant with decreased alanine levels in the *no* RPE group, implied that RPE removal switched retinal metabolism from Krebs-cycle/OXPHOS to aerobic glycolysis.

### Photoreceptors use the Krebs-cycle to produce GTP

Since in Figure 2, the *no* RPE and FCCP groups showed similar metabolite patterns and pathway changes overall, we subjected these two experimental groups to a detailed statistical analysis. This revealed 17 significant metabolite changes (Figure 4A; Supplementary Figure 7). Among the most highly reduced metabolite concentrations in the FCCP group was GTP, which positively correlated with GABA, ATP, GSSG, hypotaurine, and NAD (Figure 4B). In contrast, GDP, BCAAs, glutamine, and citrate negatively correlated to GTP. The KEGG-based pathway analysis ranked glycine, serine and threonine metabolism, glutathione metabolism, and alanine, aspartate and glutamate metabolism as the most significantly changed between the two conditions (Figure 4C).

**Figure 4.**
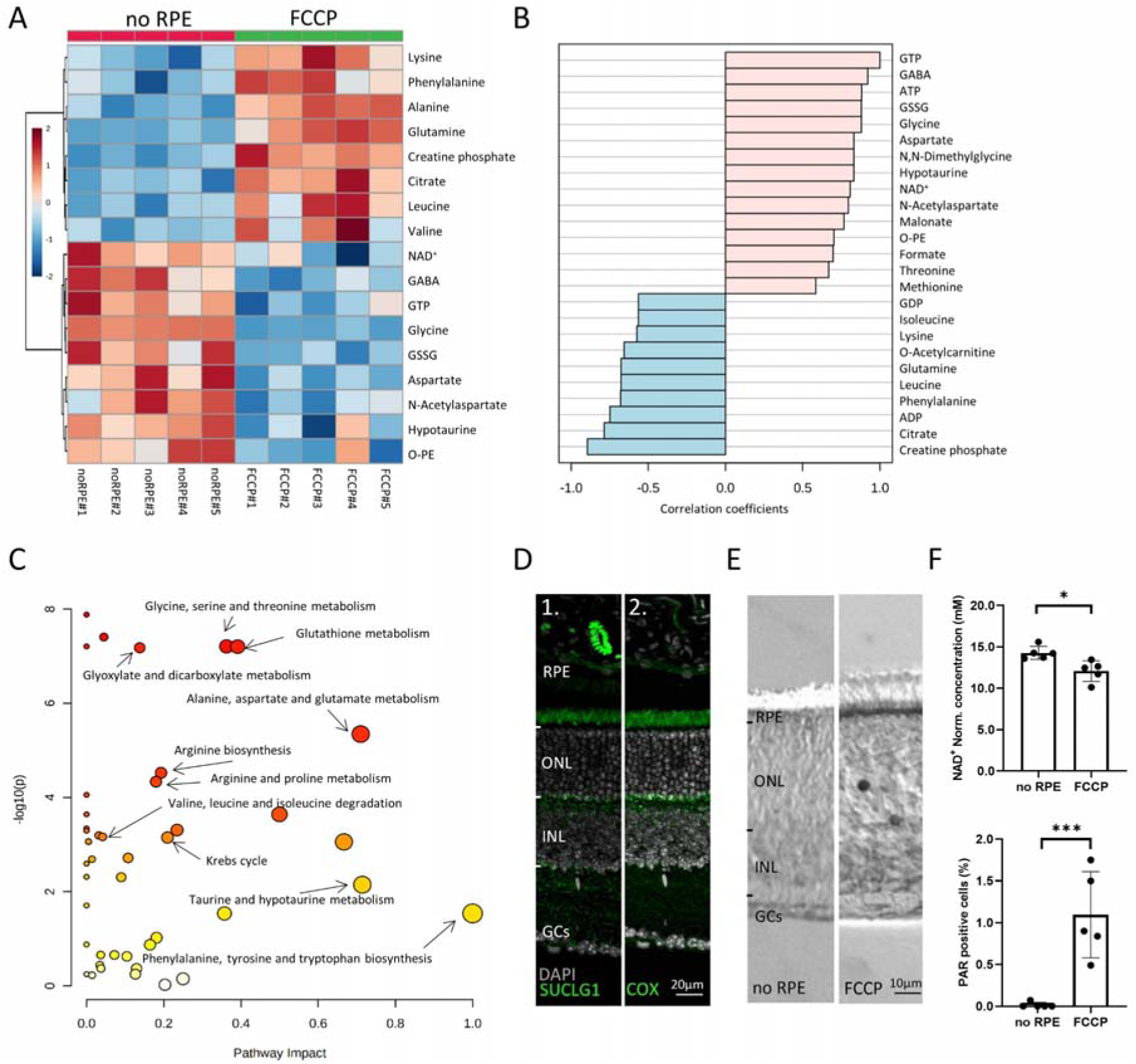
Comparison between *no* RPE and FCCP conditions. (A) Heatmap, based on unsupervised hierarchical cluster analysis by Ward’s linkage, illustrating statistically significant metabolite changes (unpaired *t*-test, fold change > 1.2, raw *p*-value < 0.05). (B) Pattern hunter for guanosine triphosphate (GTP) showing the top 25 correlating compounds. Correlation coefficient given as Pearson r distance. (C) KEGG-based pathway analysis, comparison between *no* RPE and FCCP. (D) Immunofluorescence for succinate-CoA ligase-1 (SUCLG1, green) labelled photoreceptor inner segments and colocalized with COX. DAPI (grey) was used as a nuclear counterstain. (E) PAR positive photoreceptors (black) in the outer nuclear layer (ONL) of FCCP treated retina. (F) Compared to *no* RPE, NAD^+^ levels were lower in the FCCP group, while the percentage of PAR positive cells was higher. Data represented as mean ± SD. Statistical testing: Student’s two-tailed *t*-test. *p*-values: *** < 0.001, * < 0.05. See also Supplementary Figure 7.

Curiously, measurable amounts of GTP can only be generated in two ways: 1) by succinate-CoA ligase, GTP-forming-1 (SUCLG-1) in the Krebs-cycle, and 2) from excess ATP by nucleoside-diphosphate kinase (NDK) an enzyme expressed in photoreceptor inner and outer segments (24, 25). Since ATP levels were low in both the FCCP and the *no* RPE situation, here, GTP could not have been produced from ATP. Immunofluorescence for SUCLG-1 showed strong expression in photoreceptor inner segments and synapses, where it co-localized to a large extent with mitochondrial COX (Figure 4D). Hence, the SUCLG-1 retinal expression pattern and the synthesis of GTP in the absence of RPE provided clear evidence for Krebs-cycle activity in photoreceptors.

We also identified an FCCP-induced upregulation of citrate, concomitant with a downregulation of NAD^+^. This indicated a Krebs-cycle interruption at the level of D-isocitrate to α-ketoglutarate conversion, *i.e.*, a step that requires NAD^+^. NAD^+^ is known to be consumed by poly-ADP-ribose-polymerase (PARP), which is activated by oxidative DNA damage (26) and up-regulated in dying photoreceptors (27). Therefore, we quantified poly-ADP-ribose (PAR) accumulation in retinal cells as a marker for PARP activity. Compared to the *no* RPE group, more photoreceptor cells in the FCCP-treated retina showed PAR accumulation, correlating with decreased retinal NAD^+^ levels (Figure 4E, F). Hence, FCCP-induced oxidative stress may cause increased PARP activity and decreased NAD^+^ levels, eventually interrupting the Krebs-cycle, as evidenced by citrate accumulation.

Overall, the data from the *no* RPE to FCCP group comparison showed that disruption of OXPHOS led to AA accumulation, including lysine, phenylalanine, glutamine, leucine, and valine. Notably, alanine accumulation in the FCCP group was likely caused by ALT-dependent pyruvate transamination. By contrast, metabolites high in the *no RPE* group but low with FCCP treatment were probably related to Krebs-cycle in the neuroretina. This concerned especially GTP, aspartate, and NAA.

### Reduced retinal glucose uptake promotes anaplerotic metabolism

Since GLUT1 was predominantly expressed in the RPE, we initially assumed that the metabolic response to 1,9-DDF treatment might resemble that of the *no* RPE situation. However, the patterns found between these two groups were very different (*cf.* Figure 2) and to explore these differences further, we performed an additional two-way comparison. This analysis revealed 28 significantly changed metabolites (Figure 5A; Supplementary Figure 8). When compared to the *no* RPE, the 1,9-DDF group showed changes in mitochondria and Krebs-cycle-associated metabolites (*e.g.*, high ATP, high formate, low BCAA). Conversely, the Cahill-cycle product alanine and NAA were upregulated by 1,9-DDF treatment. The pattern hunter analysis showed that metabolites formate, glutamate, taurine, and citrate were positively correlated with ATP, while, AMP, GMP, and BCAA were negatively correlated with ATP (Figure 5B).

**Figure 5.**
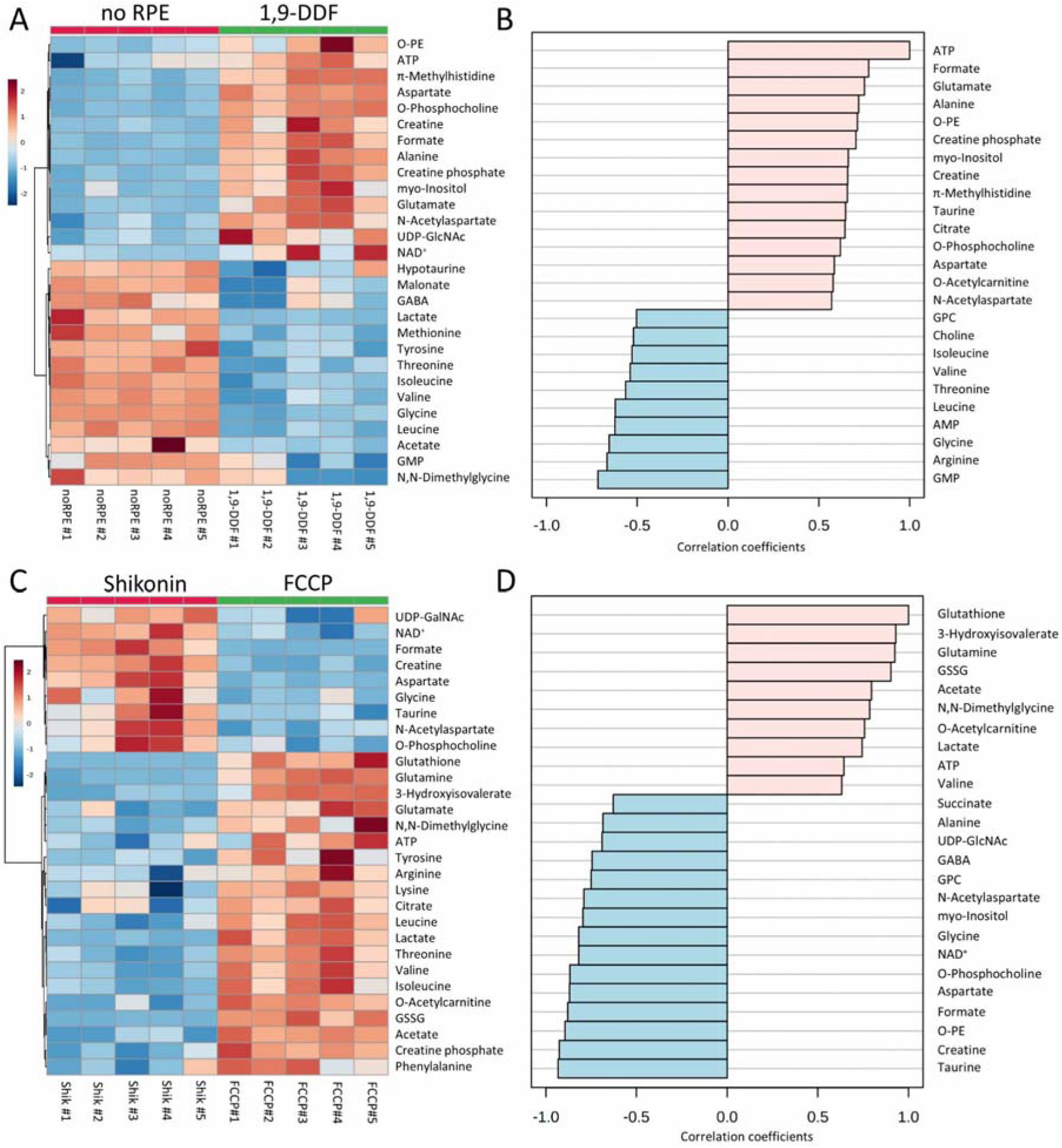
Metabolomic analysis of *no* RPE vs. 1,9-DDF treatment and Shikonin vs. FCCP treatment. Heatmap illustrating statistically significant metabolite changes (unpaired *t*-test, fold change (FC) > 1.2, raw p value < 0.05) for (A) *no* RPE vs. 1,9-DDF and (C) Shikonin vs. FCCP. Pattern hunter for (B) ATP and (D) glutathione, showing the top 25 correlating compounds. Correlation coefficient given as Pearson r distance. See also Supplementary Figures 8 and 9.

The KEGG pathway analysis showed alanine, aspartate, and glutamate metabolism, glutathione metabolism and glycine, serine, and threonine metabolism among the three most affected pathways (Supplementary Figure 8B). Taken together, the depletion of BCAAs in the 1,9-DDF treatment indicated that either the RPE, the neuroretina, or both used anaplerotic substrates to fuel the Krebs-cycle and maintain ATP production.

### Inhibition of glycolysis strongly impacts retinal ATP production

Next, we compared the Shikonin treatment, which inhibits the last step of glycolysis, to the FCCP treatment, which abolishes mitochondrial ATP synthesis, *i.e.* we compared two experimental situations that could be seen as manipulations of the two opposing ends of energy metabolism, non-oxidative glycolysis and OXPHOS. This analysis showed increased formate, aspartate, and NAA levels in the Shikonin group (Figure 5C). Conversely, higher amounts of GSH, GSSG, BCAA, threonine, and phenylalanine after FCCP treatment indicated Krebs-cycle interruption (Supplementary Figure 9). The production of lactate was higher with FCCP-compared to Shikonin treatment, probably reflecting increased glycolysis to compensate for the loss of Krebs-cycle-dependent ATP production. Metabolites connected to oxidative stress, such as GSSG and GSH, were also increased by FCCP treatment. Pattern hunter for metabolites correlated with glutathione found 3-hydroxyisovalerate, glutamine, GSSG, but also lactate and ATP, while succinate, alanine, and GABA were negatively correlated with GSH (Figure 5D). The subsequent pathway analysis identified arginine biosynthesis, alanine, aspartate, and glutamate metabolism, as well as glycolysis and gluconeogenesis as the most strongly regulated metabolic pathways (Supplementary Figure 9B).

### Blocking glucose uptake and OXPHOS disruption reveal distinct rod and cone metabolism

The metabolic rate of cones is at least two times higher than that of rods (9). The treatment with 1,9-DDF resulted in near complete cone loss, while cones were fully preserved with FCCP treatment (*cf.* Figure 1). Although we did not expect these differential treatment effects on cone survival, we used this outcome in an explorative approach to try and investigate the relative contributions of rods and cones to the metabolite patterns observed.

In this three-way comparison, 29 metabolite concentrations were significantly changed (Figure 6A; Supplementary Figure 10). The FCCP treatment reduced ATP and GTP levels, while lactate production was greatly increased, along with methionine and threonine. BCAAs, glutamine, phenylalanine and tyrosine also showed a significant upregulation with FCCP treatment. Interestingly, some metabolites like GSSG, serine, glutamate, alanine, NAD, and taurine displayed a pronounced increase in the 1,9-DDF group when compared to either control or FCCP. In addition, formate, aspartate, and myo-inositol were upregulated with 1,9-DDF treatment, together with NAA and creatine. Finally, the comparison between the 1,9-DDF and FCCP groups revealed highly increased levels of ATP and GTP, implying a strong activation of the Krebs-cycle under 1,9-DDF treatment.

**Figure 6.**
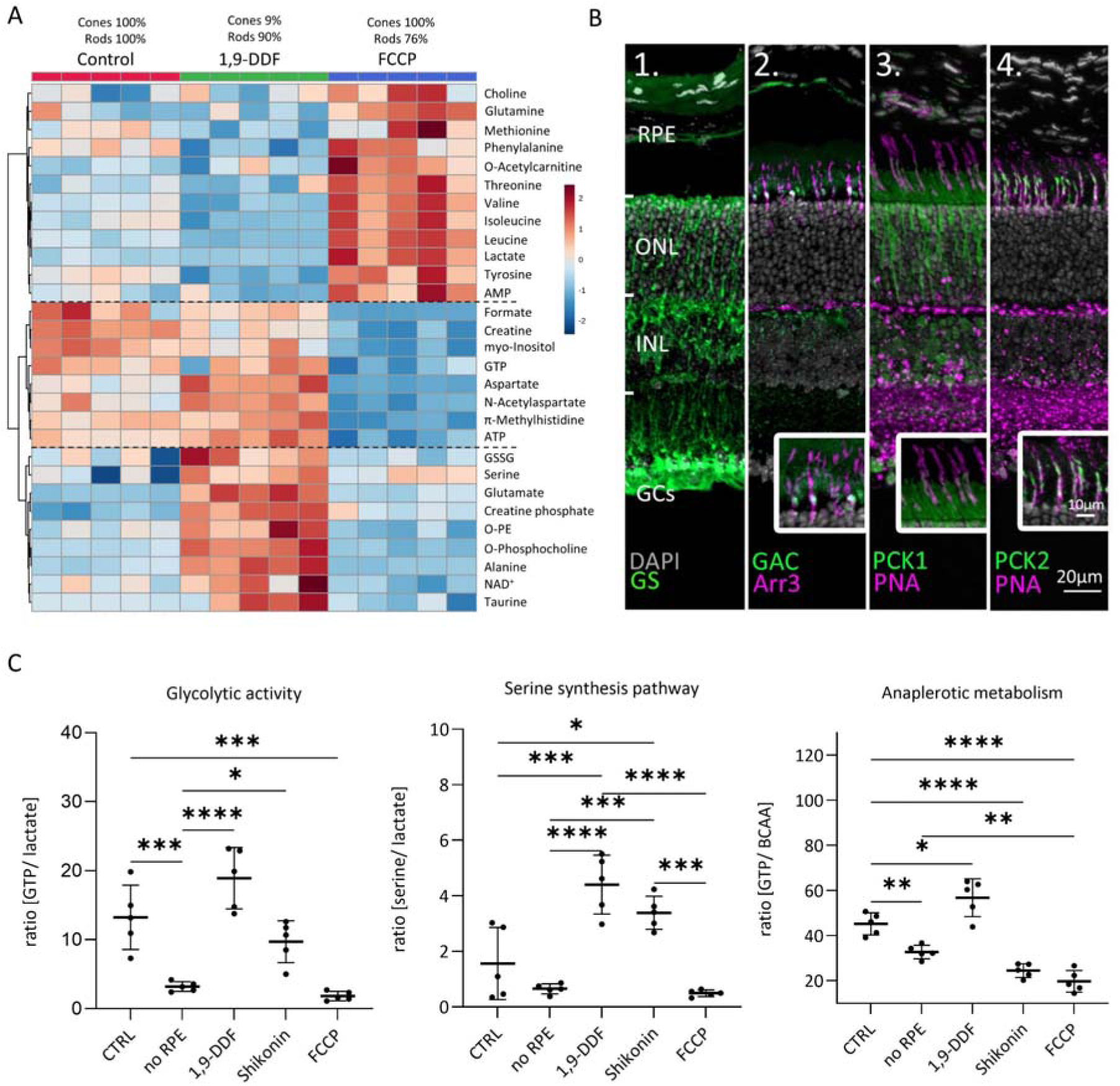
Metabolomic comparison between control, 1,9-DDF, and FCCP treatment. (A) Heatmap illustrating 29 statistically significant metabolite changes (parametric one-way ANOVA, Fisher’s LSD post-hoc analysis). Three main clusters of metabolite changes were evident (dashed lines). (B) Immunostaining (green) for enzymes related to glutamine metabolism. DAPI (grey) was used as nuclear counterstain. (B1) Glutamine synthase (GS), (B2) glutaminase C (GAC); co-localization with cone-arrestin (Arr3; magenta). (B3) phosphoenolpyruvate carboxykinase 1 (PCK1) did not co-localize with cone marker peanut-agglutinin (PNA) while (B4) PCK2 did. (C) Ratios between metabolites representing glycolysis (lactate vs. GTP), anaplerotic metabolism (GTP vs. BCAA), and serine synthesis pathway (serine vs. lactate). Data represented as individual data points with mean ± SD. Statistical testing: One-way ANOVA with Tukey’s post-hoc test; *p-*values: **** < 0.0001, *** < 0.001, ** < 0.01 * < 0.05. See also Supplementary Figure 10.

A cell type-specific attribution of these metabolite patterns may be superseded by direct drug treatment effects. Using only metabolite changes in opposing directions from control to interpret different cellular compositions, we identified BCAAs, tyrosine, NAA, and ATP (Supplementary Figure 10). BCAAs and tyrosine (and glutamine) exhibited low levels in the rod-rich 1,9-DDF group and high levels in the cone-rich FCCP group, indicating that these metabolites might be consumed by rods or produced in cones. Vice versa, it may be assumed that aspartate and NAA were predominantly produced in rods.

To investigate the role of glutamine in retinal metabolism, we used immunofluorescence (Figure 6B). We confirmed a strong expression of glutamine synthase (GS) in MGCs (28). Remarkably, the enzyme that hydrolyses glutamine to glutamate, glutaminase C (GAC) was prominently expressed in photoreceptor inner segments, photoreceptor synapses, and INL. Co-labelling with cone Arr3 revealed a strong expression of GAC in cone inner segments, implying that cones could use glutamine as a metabolic substrate. Accordingly, treatment with the GAC-specific inhibitor Compound 968 dose-dependently reduced photoreceptor viability – affecting cones more strongly than rods – while leaving the inner retina mostly intact (Supplementary Figure 6C, D).

### Impact on glycolytic activity, serine synthesis pathway, and anaplerotic metabolism

To assess how the various experimental interventions affected retinal metabolism, we calculated the ratios for pathway-specific metabolites (Figure 6C). GTP, when produced by SUCLG-1, is a marker for Krebs-cycle activity, while lactate indicates glycolytic activity. In the *no* RPE group, the GTP-to-lactate ratio was significantly lower than in control, indicating 4.1-times higher glycolytic activity. While in the 1,9-DDF and Shikonin group the GTP-to-lactate ratios were not significantly different from control, with FCCP treatment this ratio dropped to the lowest level, in line with a strong downregulation of Krebs-cycle and concomitant upregulation of glycolysis.

We then calculated the ratio of serine to lactate as an indicator for serine synthesis pathway activity. Under control conditions, and in the *no* RPE and FCCP groups, this activity was rather low, while it was strongly increased by 1,9-DDF and Shikonin treatment. In the Shikonin group, serine production from 3-phosphoglycerate and subsequent deamination to pyruvate may bypass the PKM2 block. Serine is also a precursor of phosphatidylserine, one of the three main cell membrane components. Together with a reduction of choline and high concentrations of myo-inositol, O-phosphocholine, and O-phosphoethanolamine, this may reflect high cell membrane synthesis under 1,9-DDF treatment.

Finally, we used the ratio of GTP to BCAAs to investigate to what extent Krebs-cycle activity was driven by anaplerotic metabolism. A comparatively low GTP to BCAA ratio in the *no* RPE situation showed that much of the GTP generated in the neuroretina likely came from anaplerotic substrates. While both Shikonin and FCCP treatment decreased BCAA use, 1,9-DDF treatment increased anaplerotic metabolism.

### Evidence for mini-Krebs-cycle activity

Comparing the ratios of GTP to lactate with GTP to BCAA in the *no* RPE situation showed that a large proportion of the GTP produced in the neuroretina by SUCLG-1 was derived from anaplerotic substrates rather than from pyruvate. These substrates enter the Krebs-cycle mostly at the level of α-ketoglutarate, suggesting that the neuro-retinal Krebs-cycle might not start with citrate. Instead, only an α-ketoglutarate to oxaloacetate shunt would be employed, including the GTP-synthesizing step from succinyl-CoA to succinate (Figure 7A). A key step of this mini-Krebs-cycle (28) is the transamination of oxaloacetate with glutamate by aspartate aminotransferase (AST) to yield α-ketoglutarate and aspartate, which is further acetylated to NAA. To investigate this possibility, we analyzed eight metabolites associated with the Krebs-cycle (citrate, succinate, fumarate, GTP) and the hypothesized mini-Krebs-cycle (glutamine, glutamate, aspartate, NAA).

**Figure 7.**
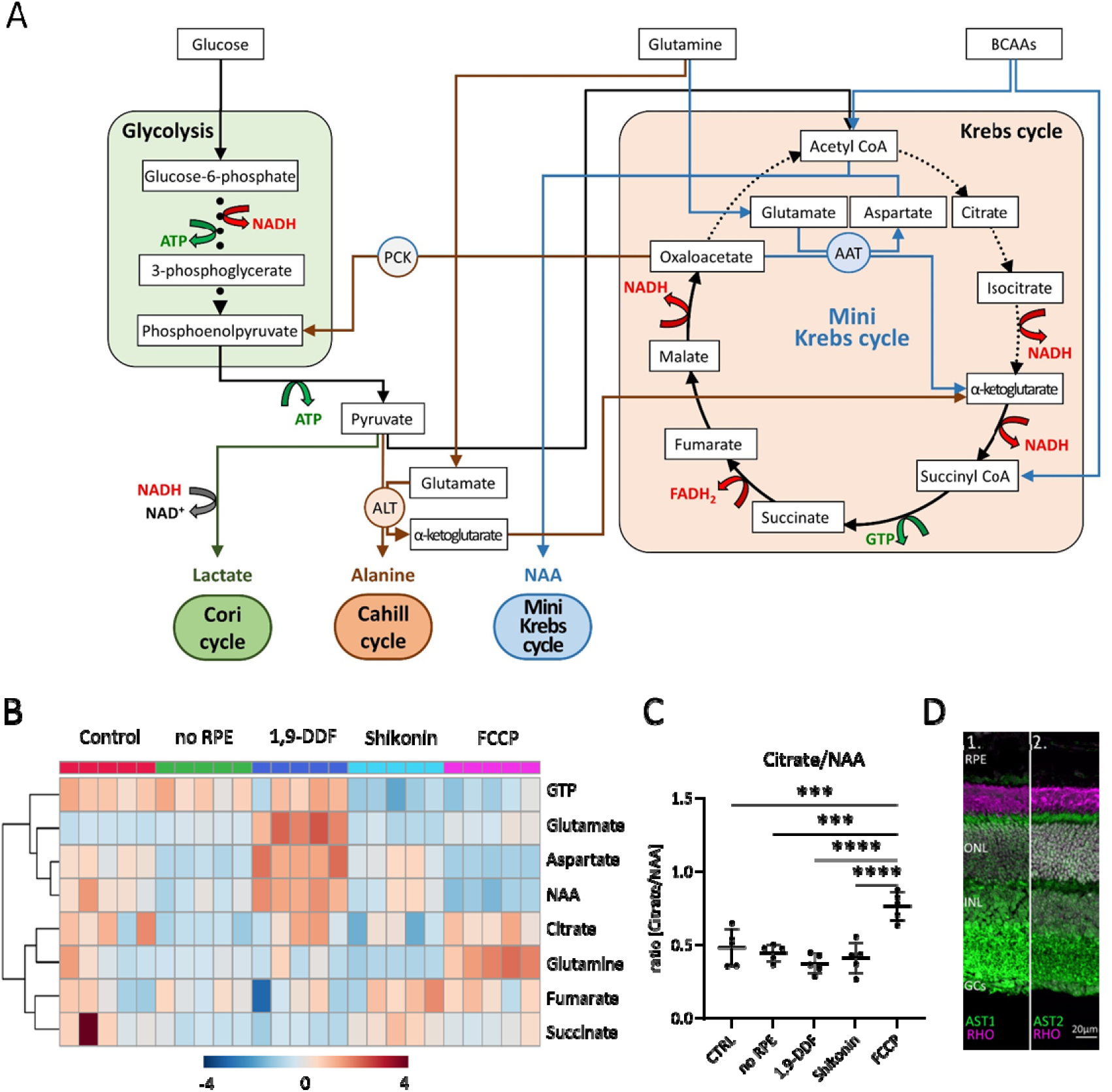
Metabolic pathways in the retina, key metabolites, and expression of aspartate amino transferase (AST). (A) Overview of main metabolic pathways and metabolites. Execution of Cori-(green arrows), Cahill-(brown), or mini-Krebs-cycle (blue) releases the signature metabolites lactate, alanine, and N-acetylaspartate (NAA), respectively. Key enzymes of the Cahill- and mini-Krebs-cycle are phosphoenolpyruvate carboxykinase (PCK), pyruvate kinase M (PKM), alanine transaminase (ALT) and AST. (B) Hierarchical clustering of eight metabolites connected to Krebs and mini-Krebs cycle. (C) Ratio NAA vs. citrate, representing full and mini-Krebs-cycle. Data represented as ratio of individual data points with mean ± SD. Statistical comparison using one-way ANOVA, Tukey’s multiple comparisons test; p-values: **** < 0.0001, *** < 0.001. (D) Immunostaining for aspartate amino transferase-1 and −2 (AST1, AST2; green) co-stained with the rod photoreceptor outer segment marker rhodopsin (RHO; magenta), DAPI (grey) was used as nuclear counterstain.

Hierarchical clustering showed similar patterns for aspartate and NAA in all experimental conditions (Figure 7B). Except for the FCCP treatment, the ratio of citrate/NAA was 0.5, indicating that the retina preferred the mini-Krebs-cycle over the full Krebs-cycle (Figure 7C). Immunolabelling for AST – conventionally associated with muscle and liver metabolism (29, 30) – found both AST1 and AST2 to be expressed in photoreceptor inner segments and cell bodies (Figure 7D). This confirmed that photoreceptors can execute the mini-Krebs-cycle, while the NAA production seen in the metabolomic data suggested that this cycle was indeed used.

Finally, we compared the energetic efficiencies of glycolysis, Krebs, Cori, Cahill, and mini-Krebs cycle (Figure 8). Using anaplerotic substrates the Cahill- and mini-Krebs cycles allow for far more efficient energy production compared to the Cori-cycle. Moreover, mitochondrial PCK2 can regenerate pyruvate from oxaloacetate. Assuming that one mole of NADH/FADH_2_ can be used to generate three moles of ATP via OXPHOS, and depending on the exact stoichiometry of glutamate/BCAA input, the mini-Krebs-cycle may deliver up to 18 moles of ATP and 2 moles of GTP per 3 carbon unit. Astonishingly, this exceeds the 15 moles of ATP and 1 mole of GTP generated in the “full” Krebs-cycle.

**Figure 8.**
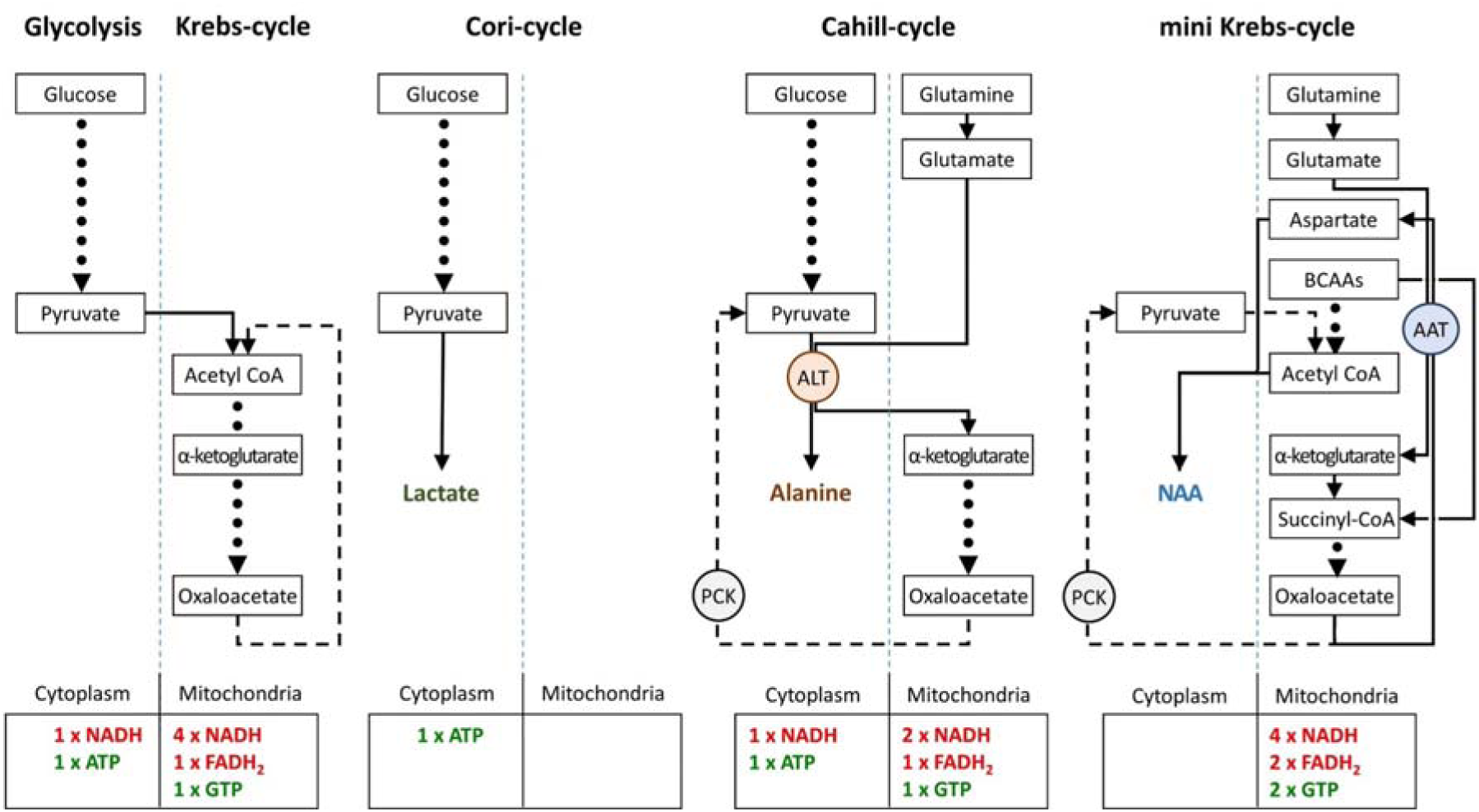
Comparison of metabolic pathways and their energetic efficiencies. Compared to the Cori-cycle, both Cahill- and mini-Krebs-cycles are highly efficient. Their key enzymes – alanine transaminase (ALT) and aspartate amino transferase (AST), respectively – generate alanine and aspartate/N-acetylaspartate (NAA). Pyruvate carboxy kinase (PCK), either in cytoplasm or within mitochondria, may reconstitute pyruvate from oxalacetate. Note that energy output of each pathway was calculated based on input of pyruvate (three carbons) or glutamate/branched-chain-amin-acid/acetyl-CoA (three carbons).

## Discussion

This study combined cellular enzyme expression patterns with quantitative metabolomics of retinal explants. In an entirely controlled environment, this enabled a comprehensive assessment of retinal metabolism and enabled an analysis of the impact of different experimental conditions on retinal cell death and survival. As opposed to earlier works, our study highlights the crucial importance of OXPHOS and anaplerotic pathways for the maintenance and survival of rod and cone photoreceptors. Importantly, we show that photoreceptors are able to uncouple glycolysis and Krebs-cycle using a NAA-producing shunt. This allows the use of both pathways in parallel and may resolve a long-standing problem in neuronal energy metabolism research. Because of the ramifications for overall cellular physiology, these findings are highly relevant for future therapy developments, for retinal diseases, as well as for neurodegenerative and further metabolic diseases in general.

### The retina as an experimental system for studies into neuronal energy metabolism

The study of neuronal metabolism in a living mammalian is notoriously difficult as there is constant interaction between different subcellular compartments, cell types, tissues, and organs (31). The retina may be a special case as it can function in isolation, *e.g.* as in an explant (32), which is why many prior studies on retinal metabolism employed explanted retina. However, in most retinal explant preparations the RPE separates from the neuroretina. Such explants were used in Otto Warburǵs seminal studies in the 1920s, where the retina was found to release large amounts of lactate (4). Later studies by Barry Winkler, using a similar experimental setup namely explanted rat retina without RPE (23), confirmed Warburǵs original work and expanded on the idea that the neuroretina was using aerobic glycolysis.

*In vivo* studies on retinal metabolism are significantly more challenging and hence relatively rare. For example, in cats, the blood lactate concentration, sampled via choroidal vein cannulation, was compared against femoral artery lactate levels, concluding that a major fraction of retinal glucose was used for aerobic glycolysis (33). In contrast, monkey *in vivo* micro-electrode measurements indicated major oxygen consumption in photoreceptor inner segments, implying the use of OXPHOS (34). Future more comprehensive *in vivo* studies of retinal metabolism may require the simultaneous cannulation of central retinal artery and central retinal vein, combined with oxygen-sensing and metabolomic analysis. Such experiments may need to employ larger animal species (*e.g.* dog, monkey) to overcome limitations in eye and blood sample size.

Our *in vitro* approach may be seen as a compromise between earlier *in vitro* studies and the *in vivo* situation. Organotypic retinal explants maintain the histotypic context of the tissue, the interactions between its various cell types, full functionality and light responsiveness, and are cultured under defined conditions, devoid of serum and antibiotics (32, 35). These retinal explants are derived from early post-natal retina (day 9) and can be cultured for up to six weeks *in vitro* without major cell loss (36). Importantly, our retinal explant cultures include the RPE (except for the *no* RPE condition) and may thus be closer to the *in vivo* situation than previous *in vitro* studies. While the combination of this *in vitro* approach with ^1^H-NMR metabolomics can be very powerful, it assumes that mainly photoreceptors are responsible for changes in retinal metabolism. Given the large photoreceptor energy consumption (5, 11) this assumption may often be correct, yet caution is warranted to not miss metabolic responses of smaller cell populations (*e.g.* Müller glia, RPE cells).

### Photoreceptors may use both glucose and anaplerotic substrates as fuel

Under anaerobic conditions, the production of lactate from glucose is a hallmark of skeletal muscle function. Here, pyruvate is reduced to lactate, which enters the bloodstream to serve as substrate for liver gluconeogenesis. Glucose can then be cycled back to the muscle. This glucose-lactate cycle between muscle and liver was discovered in the 1930s and is referred to as Cori-cycle (12). The release of large amounts of lactate from isolated retina suggested that also the retina might use the Cori-cycle, albeit under aerobic conditions (37). Photoreceptors have thus been proposed to consume primarily glucose via aerobic glycolysis (11). The resultant lactate would be used as fuel in the Krebs-cycle by RPE and MGCs, generating ATP via OXPHOS. However, this hypothesis is contradicted by the high density of mitochondria in photoreceptor inner segments (38). Moreover, key enzymes for Krebs-cycle (CS, FH, SUCLG1) and OXPHOS (COX, ATP synthase γ) were strongly expressed in photoreceptors, but showed little or no expression in RPE or MGCs, in line with previous research (25). When we cultured retina without RPE, as done by Warburg and others (4, 11), we indeed confirmed a strong lactate production. Yet, in retina cultured with RPE, lactate production was minor. Moreover, retina without RPE still produced large amounts of GTP, likely stemming from SUCLG1 activity and indicating Krebs-cycle operation in photoreceptors. Neuroretina without RPE displayed reduced viability of both rod and cone photoreceptors, as well as an accumulation of BCAAs and lactate. In contrast, the block of glucose import into the RPE by treatment with 1,9-DDF (39) resulted in a depletion of BCAAs and other AAs, and an increase of retinal ATP, indicating a switch from glycolytic to anaplerotic metabolism (15). This is in line with an earlier study using a conditional GLUT1 knockout in the RPE, which yielded evidence for a compensatory upregulation of anaplerotic metabolism in photoreceptors (3). Hence, photoreceptors likely consume both glucose and anaplerotic substrates as fuels, with retinal metabolism switching to aerobic glycolysis only in the absence of RPE. This switch may be driven by increased glucose uptake.

In intact retina, tight junction-coupled RPE cells form the outer blood-retinal barrier and prevent direct access of glucose to photoreceptors (40). In the absence of RPE, photoreceptors expressing the high affinity/high capacity GLUT3 become “flooded” with glucose. The resultant Crabtree effect likely causes a shutdown of Krebs-cycle activity (18, 41, 42), effectively starving photoreceptors to death because a purely glycolytic metabolism may not generate sufficient ATP for their survival. As seen in the *no* RPE group, cones appear to be more sensitive to this effect, perhaps because their overall energy demand is twice as high as that of rods (9). Surprisingly, FCCP treatment did not reduce cone viability suggesting that for cones high glucose is more toxic than a lack of OXPHOS. At any rate, it is probably the lack of the blood-retinal barrier function that leads to the strong differences in metabolite patterns seen between the 1,9-DDF treatment and the *no* RPE group. Taken together, the high rates of retinal aerobic glycolysis first reported by Otto Warburg are likely an artefact of the absence of RPE, while intact retina may rely more on OXPHOS for its energy production.

### Rods need OXPHOS, cones need glycolysis

Treatment with FCCP revealed striking differences between rod and cone energy metabolism. FCCP eliminates the proton gradient between inner mitochondrial membrane and mitochondrial matrix, abolishing ATP synthesis (43). Initially, FCCP may increase Krebs-cycle activity to attempt restoration of mitochondrial proton gradient (44), increasing oxidative stress (45), and oxidative DNA damage. The resultant activation of PARP would deplete NAD, further aggravating metabolic stress (46). Although FCCP had a strong toxic effect on rod photoreceptors, remarkably, cone photoreceptors were almost completely preserved. Theoretically, it is conceivable that FCCP primarily affected the RPE and that photoreceptor degeneration occurred secondarily to that. However, then, rods and cones would likely have been equally affected by FCCP treatment. In striking contrast to the FCCP effect was that of the 1,9-DDF block on GLUT1 (39). Although GLUT1 was strongly expressed in the RPE, glucose entering the RPE is shuttled forward to photoreceptors (3). Remarkably, with 1,9-DDF, over 90 % of cones were lost, while the detrimental effect on rods was comparatively minor. These differential effects of FCCP and 1,9-DDF treatments strongly suggest that glycolysis is sufficient and necessary for cone survival, in agreement with earlier studies who showed that under stress cones require glycolysis to remain viable (47, 48).

Conversely, rods require OXPHOS for their survival, while glycolysis is of minor importance. Furthermore, cone viability relative to rods was also strongly compromised in the *no* RPE situation, indicating that an intact blood-retinal barrier (40) and regulation of glucose access was important for cone survival. Future studies analyzing specific mouse mutants which have either only rods (*e.g*. the *cpfl1* mouse (49) or only cones (*e.g.* the NRL knock-out mouse (50) may give more detailed insights into rod and cone metabolism.

### Expression of glucose transporters in the outer retina

Our work shows that RPE cells express high levels of GLUT1, while photoreceptors express the neuronal glucose transporter GLUT3. Furthermore, GLUT2 may contribute to photoreceptor glucose uptake indirectly via horizontal cells (51). We note that there is a controversy in the literature regarding the expression of GLUT1 on photoreceptors. Although there seems to be a general agreement that GLUT1 is expressed on RPE cells (3), several studies have suggested GLUT1 expression also on photoreceptors, both on rods and cones (48, 52) In the brain the generally accepted setup for GLUT1 and GLUT3 expression is that low-affinity GLUT1 (Km = 6.9 mM) is expressed on glial cells, which contact blood vessels, while high-affinity GLUT3 (Km = 1.8 mM) is expressed on neurons (53, 54). This setup matches decreasing glucose concentration with increasing transporter affinity, for an efficient transport of glucose from blood vessels to glial cells to neurons. In the retina, the cells that contact the choroidal blood vessels are the tight-junction-coupled RPE cells. Since RPE cells express GLUT1, an efficient glucose transport from the RPE to the energy-hungry photoreceptors, requires photoreceptors to express a glucose transporter with a glucose affinity higher than GLUT1. Our finding of GLUT3 expression on photoreceptors matches this requirement. Still, at this point we cannot exclude a low level expression of GLUT1 on other retinal cell types, for instance on Müller glia cells (55) which might use GLUT1 to export excess glucose resulting from gluconeogenesis (56) Moreover, there are 10 further glucose transporters, some of which might be expressed in the retina. A comprehensive study of the retinal protein expression of all glucose transporters, using, for example, CODEX technology (57), may reveal their expression patterns and putative functions in the future.

### Cone photoreceptors likely employ the Cahill-cycle

The lactate-generating Cori-cycle (12) is highly inefficient and in intact retina likely plays a lesser role. An efficient pathway is the glucose-alanine cycle, or Cahill-cycle, in which pyruvate, instead of being reduced to lactate, is transaminated to alanine (14). This preserves NADH and generates α-keto acids to fuel the Krebs-cycle. The alanine generated enters the bloodstream and is taken up by the liver, where the ammonia is excreted in the form of urea, while the carbon backbone is used for gluconeogenesis. The key enzyme for the Cahill-cycle is ALT, conventionally associated with muscle and liver function (29, 30). Previously, ALT was found in glial cells of the honeybee retina (44), and ALT activity was detected in rat retinal tissue lysates (58). In our study, the localization of ALT in photoreceptors and inner retinal neurons, combined with our interventional and metabolomic datasets, provided evidence for the operation of the Cahill-cycle in the mammalian retina. Moreover, the expression of mitochondrial PCK2 in cones facilitates the efficient uncoupling of glycolysis from the Krebs-cycle, suggesting that cones may use the Cahill-cycle for very effective energy production.

### Glutamine from MGCs may provide additional fuel for cones

MGCs are known for their uptake of extracellular glutamate and use for glutamine synthesis. In fact, in retinal histology, glutamate-aspartate transporter (GLAST) and glutamine synthase (GS) have been widely used as markers for MGCs (25, 28). Glutaminase C (GAC) converts glutamine back to glutamate, which could then serve as a substrate for the mini-Krebs-cycle. We localized GAC in inner retinal neurons and photoreceptors, with a particularly strong expression in cone inner segments. Here, GAC overlapped with ALT expression, indicating that cones can use glutaminolysis to obtain extra glutamate for pyruvate transamination in the Cahill-cycle. This may also explain why the glycolysis inhibitor Shikonin reduced cone viability much less than that of rods. In this situation, cones might be able to use the serine-synthesis pathway – with serine production from 3-phosphoglycerate and subsequent deamination to pyruvate – to bypass the Shikonin block of PKM2.

### A hypotaurine-taurine shuttle can transfer NADH from RPE to photoreceptors

The regeneration of the photoreceptor photopigment retinal is performed by the RPE (16). Recently, retinal has been proposed to form a Schiff base adduct with taurine, which would act as a retinal carrier and buffer (59). Moreover, taurine has been suggested to be an important osmolyte in the retina that may flow in and out of neuronal cells in response to neuronal activity (60, 61).

We found that retina cultured with RPE harbored high levels of taurine and low levels of hypotaurine, while retina without RPE displayed low levels of taurine and high levels of hypotaurine. Our taurine immunostaining found essentially no taurine in the RPE, while photoreceptor inner segments and synapses displayed very high taurine levels. Together, these findings suggest a hypotaurine-taurine shuttle between RPE and photoreceptors. In the RPE taurine can be reduced to hypotaurine and shuttled back to photoreceptors where oxidation by hypotaurine dehydrogenase (59, 62) may reconstitute taurine, yielding additional NADH for OXPHOS. The net effect of this hypotaurine-taurine shuttle would be a transfer of reducing power from RPE to photoreceptors, boosting photoreceptor ATP production via OXPHOS. In addition, it seems likely that hypotaurine-taurine shuttling will have an impact on the osmolarity of both photoreceptors and RPE cells.

### Rods can use the mini-Krebs-cycle to uncouple glycolysis from mitochondrial metabolism

A key problem in the understanding of cellular energy metabolism is the relationship between fast but inefficient glycolysis and slow but efficient Krebs-cycle/OXPHOS (63, 64). Both pathways are coupled via pyruvate and the different metabolic flow rates reduce the efficiency of energy production, *e.g.*, via feedback inhibition (65). Pyruvate coupling is especially problematic in high and rapidly changing energy demand such as in neurons and photoreceptors (66). Uncoupling glycolysis and Krebs-cycle via the Cori-cycle is extremely wasteful and likely insufficient to satisfy long-term photoreceptor energy demand. By comparison, the Cahill-cycle delivers additional NADH when using pyruvate derived from glycolysis, and cones may use the Cahill-cycle for uncoupling from glycolysis.

An alternative pathway is the mini-Krebs-cycle, essentially an oxalacetate to α-ketoglutarate shunt (15). This cycle uses glutamate, glutamine, and BCAAs as fuels to run mitochondrial respiration independent of glycolysis. The key step of the mini-Krebs-cycle is the transamination of oxaloacetate/glutamate to aspartate/α-ketoglutarate by AST. α-Ketoglutarate is then metabolized to oxaloacetate, generating NADH, FADH_2_, and GTP. Acetyl-CoA generated from BCAAs or pyruvate is used to create NAA, the end product of the mini-Krebs-cycle. We found mitochondrial AST2 to be expressed in rod inner segments, in agreement with an early cytochemical study (67). This indicates that the mini-Krebs-cycle is used primarily by rods, perhaps explaining their selective vulnerability to FCCP treatment. Crucially, the mini-Krebs-cycle is more energy-efficient than the Cahill-cycle and generates NAA instead of alanine as net product. Both metabolites serve the purpose of disposing of excess ammonia originating from AA input (68, 69). NAA in the human brain is one of the most abundant metabolites and is routinely used in clinical MRI diagnosis to visualize brain health (70). While different functions have been hypothesized for NAA (71, 72), our work proposes NAA as a signature metabolite for the mini-Krebs-cycle. Indeed, a recent study using Raman spectroscopy imaging detected high levels of NAA in the human retina, demonstrating *in vivo* use of this cycle (73). Importantly, the 6-step mini-Krebs-cycle is significantly faster than the 10-step Krebs-cycle (15), more energy-efficient, and uncoupled from glycolysis. While the high metabolic rate of photoreceptors suggests that mini-Krebs-cycle metabolites will mostly originate there, the strong AST1/AST2 expression observed in the inner plexiform layer suggests that the mini-Krebs-cycle may also play a role in fueling synaptic activity. Moreover, in rods, PCK1 may allow to replenish cytoplasmic acetyl-CoA pools from oxaloacetate, if BCAA-derived input was insufficient.

In the future, it will be very interesting to perform carbon- or nitrogen tracing studies using C-or N-labelled alternative fuels, including glucose, lactate, and glutamate/glutamine, to measure which metabolites are produced under what experimental conditions. Such tracing studies would also allow to establish the flux of metabolites through a specific pathway, something that is difficult to assess with our current steady-state measurements.

Overall, the different pathways outlined here provide photoreceptor cells with remarkable versatility and flexibility in terms of substrate use, enabling them to dynamically adapt the timing and quantitative output of energy metabolism. Rods rely strongly on the Krebs-cycle and OXPHOS, while cones are more dependent on glycolysis. These differences between rod and cone metabolism may be related to their response kinetics and sensitivities. The flexible uncoupling of glycolysis from mitochondrial respiration, allows both processes to run at optimum, producing the characteristic signature metabolites lactate (Cori-cycle), alanine (Cahill-cycle), and NAA (mini-Krebs-cycle). These metabolites can reveal energy or disease status and could serve as readout and guide for the design of novel therapeutic interventions. Given the general importance of energy metabolism, the significance of our findings extends beyond the retina, for instance, to other neurodegenerative and metabolic diseases.

## Materials and Methods

C3H wild-type (WT) mice were used to generate organotypic retinal explant cultures (18). All efforts were made to minimize the number of animals used and their suffering. For details on the experimental procedures, immunofluorescence, and antibodies, as well as metabolomic analysis and databases used please refer to the supporting information (SI).

## Declarations

## Ethics approval and consent to participate

Protocols compliant with the German law on animal protection were reviewed and approved by the institution for animal welfare, veterinary service and laboratory animal science (Einrichtung für Tierschutz, Tierärztlichen Dienst und Labortierkunde) of the University of Tübingen and were following the association for research in vision and ophthalmology (ARVO) statement for the use of animals in vision research.

## Availability of data and materials

Raw NMR spectroscopy data is available upon request. Any additional information required to reanalyze the data reported in this paper is available from the lead contact upon request. This paper does not report original code.

## Competing interests

The authors declare no competing financial interests. CT reports a research grant by Bruker BioSpin GmbH.

## Funding

This work was funded by the ProRetina Foundation, the Zinke heritage foundation, the Werner Siemens Foundation, the Chinese scholarship council (CSC), ANID-FONDECYT No. 1210790 (OS) and PhD grant BECAS CHILE/2018 −21180443 (VC). We also acknowledge support from the Open Access Publication Fund of the University of Tübingen.

## CRediT author contribution statement

**Yiyi Chen** : Conceptualization, Methodology, Investigation, Writing – original draft. **Laimdota Zizmare** : Conceptualization, Methodology, Investigation, Writing – original draft, Writing – Review & Editing. **François Paquet-Durand**: Conceptualization, Methodology, Investigation, Writing – original draft, Writing – Review & Editing, Supervision, Funding acquisition. **Christoph Trautwein** : Conceptualization, Methodology, Investigation, Writing – original draft, Writing – Review & Editing, Supervision, Funding acquisition. **Shirley Yu**: Investigation. **Lan Wang**: Investigation. **Friedrich W. Herberg**: Writing – Review & Editing. **Oliver Schmachtenberg**: Writing – Review & Editing. **Victor Calbiague**: Writing – Review & Editing.

## Acknowledgements

We thank N. Rieger, M. Owczorz, and D. Bucci for first-rate technical support, as well as James B. Hurley and Daniel Hass (both University of Washington, Seattle, USA) for helpful comments and suggestions.

## Abbreviations

1,9-DDF: 1,9 dideoxyforskolin
AA: amino acids
AST: aspartate amino transferase
ADP: adenosine diphosphate
ALT: alanine transaminase
ATP: adenosine triphosphate
BCAA: branched chain amino acid
COX: cytochrome oxidase
GAC: glutaminase C
GCs: ganglion cells
GLUT: glucose transporter
GS: glutamine synthase
GTP: guanosine triphosphate
GABA: gamma amino butyric acid
GPC: sn-glycero-3-phosphate
FCCP: carbonyl cyanide-*p*-trifluoromethoxyphenylhydrazone
GSSG: glutathione disulfide
INL: inner nuclear layer
MGC: Müller glial cells
NAA: N-acetylaspartate
NAD: nicotinamide adenine dinucleotide
ONL: outer nuclear layer
O-PE: O-phosphoethanolamine
OXPHOS: oxidative phosphorylation
PARP: poly(ADP)ribose polymerase
PCK: phosphoenolpyruvate carboxy kinase
PKM: pyruvate kinase M
PNA: peanut agglutinin
RP: retinitis pigmentosa
RPE: retinal pigment epithelium
RPE65: retinal pigment epithelium-specific 65 kDa protein
SUCLG1: succinate-CoA ligase-1
TUNEL: terminal UDP nick-end labelling
UDP: uracil diphosphate

## Notes

### Competing Interest Statement

The authors have declared no competing interest.

### Summary of Updates

format updated

