## Supplementary material for "Retinal metabolism: Evidence for uncoupling of glycolysis and oxidative phosphorylation via Cori-, Cahill-, and mini-Krebs-cycle": SI materials+methods

**This PDF file includes:**

Supporting Information (SI) of Materials and Methods

Supplementary Tables S1 to S2

Supplementary Figures S1 to S10

SI References

Supporting Information

**Materials and Methods**

**Animals**

C3H wild-type (WT) mice were used (1). These animal lines are regularly screened for genetic mutations known to cause retinal degeneration (*e.g. rd1, rd2, rd8, rd10, cpfl1*, *etc*.) and were shown to be free of such mutations. All efforts were made to minimize the number of animals used and their suffering. Animals were housed under standard white cyclic lighting, had free access to food and water, and were used irrespective of gender. Protocols compliant with the German law on animal protection were reviewed and approved by the ‘‘Einrichtung fur Tierschutz, Tierärztlichen Dienst und Labortierkunde’’ of the University of Tübingen and were following the association for research in vision and ophthalmology (ARVO) statement for the use of animals in vision research. Animals were not assigned to experimental groups prior to their sacrifice.

**Retinal explant cultures**

The retinal explantation procedure and long-term cultivation in defined medium, free of serum and antibiotics, is described in detail in (2) (*cf*. Supplementary Figure 1B). Briefly, mice were decapitated at postnatal day (P) 9 and the heads cleaned with 70% ethanol. The eyes were removed under aseptic conditions, and placed into R16 basal medium (BM; Gibco, Paisley, UK), washed for 5 min, followed by a 15 min incubation in 0.12% Proteinase K (Sigma-Aldrich, Taufkirchen, Germany; P6556) at 37°C to predigest the sclera and to allow for an easy separation of the retina together with its retinal pigment epithelium (RPE). Then, the eyes were placed for 5 min in BM with 10% foetal calf serum (FCS) to deactivate Proteinase K. In the case of no RPE explants, the retina was explanted directly, without Proteinase K pre-treatment and FCS deactivation. Under the microscope and sterile conditions, the anterior segment, lens, and vitreous body were carefully removed from the eyeballs, the optic nerve was cut, and the retinas were removed from the sclera. Then four incisions were made into the retina to give a flat, clover leaf like structure that was transferred to a culturing membrane (sterile 24 mm insert with 0.4 µm pore size polycarbonate membrane, Corning-Costar, New York, NY, USA), with the ganglion cell layer facing up. Subsequently, culturing membranes were placed in six-well culture plates (BD Biosciences, San Jose, CA, USA) and incubated in 1 ml of R16 complete medium (CM) with supplements (64), at 37°C, in a humidified incubator with 5% CO2. From P9 to P11 cultures were kept in CM without treatment to adapt to *in vitro* conditions. Drug treatments were applied to the culturing medium from P11 until P15 with either 50 µM 1,9-DDF (Sigma-Aldrich), 4 µM Shikonin (3-5) (Selleck Chemicals, Planegg, Germany), 20-400 µM β-chloro-alanine (Sigma-Aldrich), or 0.1-10 µM Compound 968 (Biomol, Hamburg, Germany). The FCCP treatment (5 µM) was applied from P13 to P15. This form of drug application meant that the drugs reached the RPE and neuroretina from the basal side (i.e. through the porous culturing membrane). The medium was changed every 2 days.

For all treatment conditions, culturing was stopped at P15 by 45 min fixation in 4% paraformaldehyde (PFA). Retinal tissues were cryoprotected with graded sucrose solutions containing 10, 20, and 30% sucrose and then embedded in Tissue-Tek O.C.T. compound (Sakura Finetek Europe, Alphen aan den Rijn, Netherlands). Tissue sections of 12 µm were prepared using Thermo Scientific NX50 microtome (Thermo Scientific, Waltham, MA) and thaw-mounted onto Superfrost Plus glass slides (R. Langenbrinck, Emmendingen, Germany).

**Cell death detection (TUNEL assay)**

Fixed slides were dried at 37°C for 30 min and washed in phosphate buffered saline (PBS) solution at room temperature (RT), for 15 min. Afterwards, the slides were placed in TRIS buffer with proteinase K at 37°C for 5 min to inactivate nucleases. The slides were then washed with TRIS buffer (10 mM TRIS-HCL, pH 7.4), 3 times for 5 minutes each. Subsequently, the slides were placed in ethanol-acetic acid mixture (70:30) at -20°C for 5 min followed by 3 washes in TRIS buffer and incubation in blocking solution (10% normal goat serum, 1% bovine serum albumin, 1% fish gelatine in 0.1% PBS-Triton X100) for 1h at RT. Lastly, the slides were placed in the terminal dUTP-nick-end labelling (TUNEL) solution (labelling with either fluorescein or tetra-methyl-rhodamine; Roche Diagnostics GmbH, Mannheim, Germany) in 37°C for 1 h and mounted with Vectashield with DAPI (Vector, Burlingame, CA, USA) thereafter.

**Immunofluorescence**

Fixed slides were dried at 37°C for 30 min and rehydrated for 10 min in PBS at RT. For immunofluorescent labelling, the slides were incubated with blocking solution (10% normal goat serum, 1% bovine serum albumin in 0.3% PBS-Triton X 100) for 1 h at RT. The primary antibodies were diluted (see supplemental table 2) in blocking solution and incubated at 4°C overnight. The slides were then washed with PBS, 3 times for 10 min each. Subsequently, the secondary antibody, diluted in PBS (see table 1), was applied to the slides, and incubated for 1 h at RT. Lastly, the slides were washed with PBS and mounted with Vectashield with DAPI (Vector).

**Microscopy, cell counting, and statistical analysis**

Fluorescence microscopy was performed with a Z1 Apotome microscope equipped with a Zeiss Axiocam digital camera (Zeiss, Oberkochen, Germany). Images were captured using Zen software (Zeiss) and the Z-stack function (14-bit depth, 2752*2208 pixels, pixel size = 0.227 µm, 9 Z-planes at 1 µm steps). The raw images were converted into maximum intensity projections using Zen software and saved as TIFF files.

Photoreceptors stained by the TUNEL assay were counted manually on three images per explant, the average cell number in a given ONL area was estimated based on DAPI staining and used to calculate the percentage of TUNEL positive cells. Cones stained with arrestin-3 were counted on Apotome images using maximum intensity projection (MIF). Cone numbers are expressed as arrestin-3 positive cells visible in 100 µm stretches of retinal circumference.

Adobe Photoshop CS6 (Adobe Systems Inc., San Jose, CA) and Adobe Illustrator CC 2019 software was used for primary image processing. All data given represent the means and standard deviation from at least 5 different animals. Statistical comparisons between experimental groups were made using Student´s paired *t*-test or ANOVA and multiple comparisons correction with Tukey´s post-hoc test (*cf*. Figure 1) using Graph Pad Prism 9.1 for Windows (Graph Pad Software, La Jolla, CA). Levels of significance were: * = *p* < 0.05; ** = *p* < 0.01; *** = *p* < 0.001.

**Metabolite extraction**

After retinal explant culture, at P15, the tissue was quickly transferred into 80% methanol / 20% ethanol, snap-frozen in liquid nitrogen. A sample of the culture medium was taken from the same well plate as the retinal tissue, and snap-frozen in liquid nitrogen. Retinal tissue was placed in 400 µL of methanol (LC-MS grade), transferred to the 2 mL glass Covaris system-compatible tubes and 800 µL of methyl-tert-butyl ether (MTBE) was added, thoroughly mixed, and further subjected to metabolite extraction via ultra-sonication (Covaris E220 Evolution, Woburn, USA). After the extraction, 400 µL of ultrapure water were added for two-phase liquid separation. The aqueous phase was separated and evaporated to dryness. Similarly, 400 µL of the aqueous medium sample was transferred to the 2 mL glass Covaris system-compatible tube, 800 µL of MTBE was added and subjected to the ultra-sonication extraction protocol. Finally, 400 µL of methanol were added and mixed, centrifuged, and after two-phase separation the aqueous layer was separated and evaporated, to obtain a dry metabolite pellet.

**Sample preparation for ^1^H-NMR spectroscopy measurements and data analysis**

Dried metabolite pellets were resuspended in a deuterated phosphate buffer (pH corrected for 7.4) with 1 mM of 3-(trimethylsilyl) propionic-2,2,3,3-d_4_ acid sodium salt (TSP) as internal standard. NMR spectra were recorded at 298 K on a 14.1 Tesla ultra-shielded NMR spectrometer at 600 MHz proton frequency (Avance III HD, Bruker BioSpin, Ettlingen, Germany) equipped with a triple resonance 1.7 mm room temperature micro probe. Short zero-go (zg), 1D nuclear Overhauser effect spectroscopy (NOESY) and Carr-Purcell-Meiboom-Gill (CPMG; 4096 scans for retinal tissue samples, 128 for medium samples) pulse programs were used for spectra acquisition. Spectra were processed with TopSpin 3.6.1 software (Bruker BioSpin).

**Quantification of metabolomic data and statistical analysis**

Retina tissue metabolite assignment and quantification was done on the pre-processed CPMG spectra and performed with ChenomX NMR Suite 8.5 (Chenomx Inc., Edmonton, Canada). For absolute quantification, metabolite matches were calibrated to the internal 1 mM TSP standard. Of note, not all central metabolic pathways are detectable by ^1^H-NMR, which is why the fully annotated metabolite list is lacking some of the glycolytic (*e.g.* glycerylaldehyde-3-phosphate) and Krebs-cycle (*e.g.* succinyl-CoA) metabolites. After manual assignment and quantification of each sample, a raw metabolite concentration table (comma separated value) was exported and used as upload file for statistical analysis with MetaboAnalyst 5.0 platform (<https://www.metaboanalyst.ca> (6)).

Highly abundant metabolites (glucose, lactate) could be also quantified in medium samples. However, in these samples the internal standards were not well preserved due to the high salt content of the medium. Therefore, spectra were manually integrated in the TopSpin software, to obtain absolute integral values for glucose (5.2 ppm) and lactate (4.1 ppm). The glucose peak integral was assigned to the known 19.1 mM absolute glucose concentration in the unspent, complete R16 medium, and glucose and lactate concentrations in experimental samples were calculated in proportion to their absolute integrals.

Raw metabolite concentrations were normalized by the probabilistic quotient normalization (PQN) method on the control group to account for dilution effects. Heat maps display auto-scaled metabolite concentrations (red – high, blue – low). Unsupervised hierarchical cluster analysis was performed by Euclidean distance measure and Ward distance measure. The log_2_ fold change (FC) shown in Volcano plots indicates the log_2_ (mean metabolite concentration in condition A) minus log_2_ (mean metabolite concentration in condition B) where A and B refers to the two different experimental groups being compared (*e.g.* control *vs*. no RPE).

Pattern hunter correlation analysis was performed on respective datasets in each specific comparison of selected metabolites based on the most statistically significant changes and importance using Pearson r distance measure for correlation coefficient calculation. GraphPad Prism 9.1.0 software (GraphPad Software, San Diego, CA, USA) was used for statistical analysis and data visualization. For multiple group comparisons an ordinary one-way ANOVA with Tukey’s multiple comparisons post-hoc test was applied. In supplemental Figure 10 and ordinary one-way ANOVA with Fisher´s LSD test was used. For two-group comparisons Student´s unpaired, two-tailed *t*-test was used. Fold change (FC) threshold was set to > 1.2, and raw p value < 0.05, *p*-values represent: **** < 0.0001, *** < 0.001, ** < 0.01 * < 0.05. Data throughout is represented as individual data points with mean ± SD. Metabolite pathway analysis was based on Kyoto Encyclopaedia of Genes and Genomes (KEGG) pathway database where pathway impact values are based on certainty and pathway enrichment analysis.

**Data and Code availability**

Any additional information required to reanalyze the data reported in this paper is available from the corresponding authors upon request. This paper does not report original code.

**Supplementary Table S1.** Full list of all quantified metabolites in retina samples. A total of 50 metabolites were quantified in control, no RPE, 1,9-DDF, Shikonin and FCCP groups. Each metabolite mean concentration value (µM concentration range) and standard deviation (stdev) is displayed in the table (n=5).

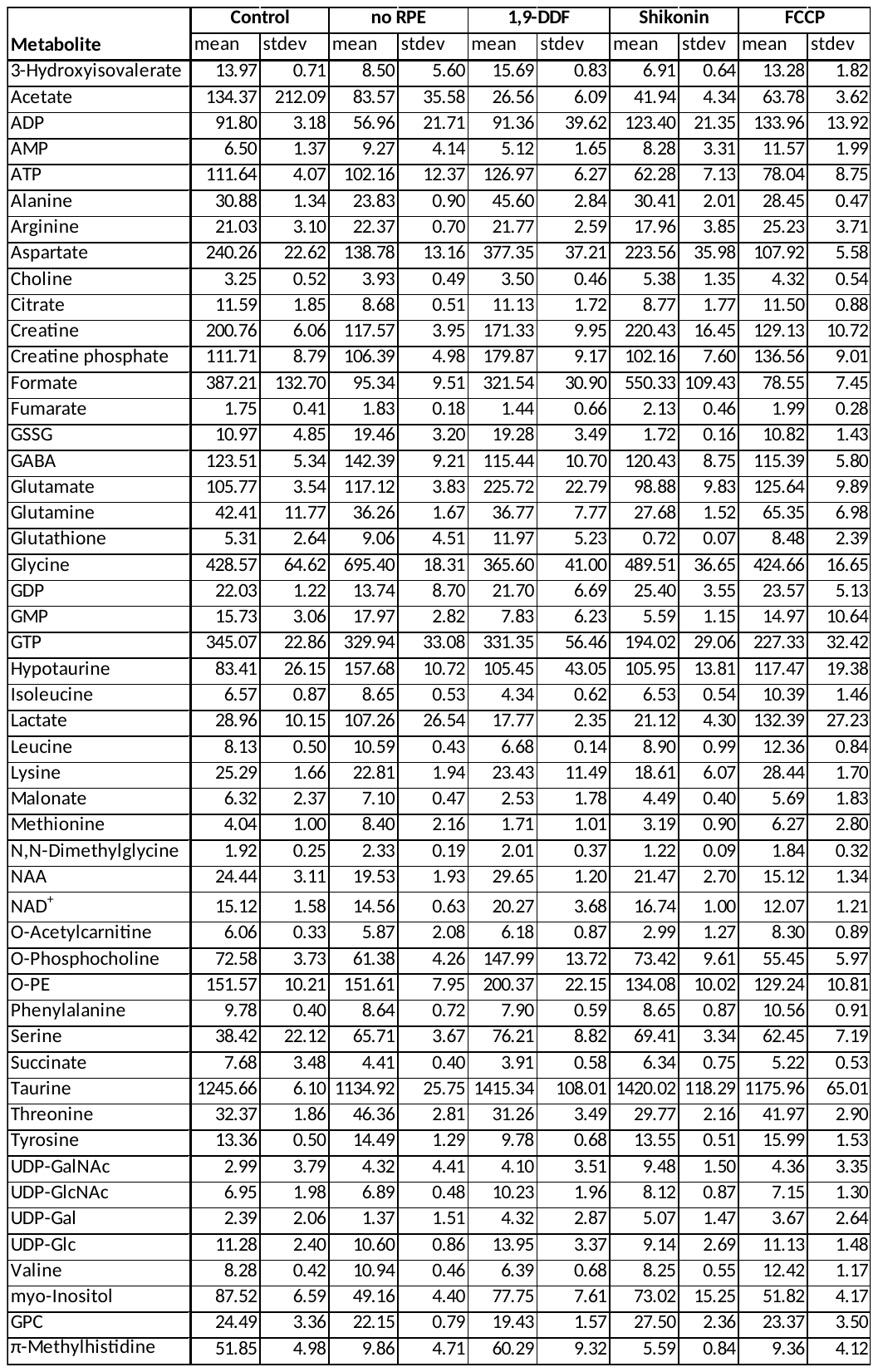

Supplementary Table S2. Primary and secondary antibodies used in the study, providers, and dilutions.

| REAGENT or RESOURCE | SOURCE | IDENTIFIER |
| --- | --- | --- |
| Antibodies, dilution | | |
| Mouse monoclonal anti-RPE65, 1:100 | Thermo Fisher Scientific | Cat# MA5-16042, RRID: AB_11151857 |
| Rabbit polyclonal anti-GLUT1, 1:300 | Abcam | Cat# ab652, RRID:AB_305540 |
| Rabbit polyclonal anti-GLUT3, 1:300 | Abcam | Cat# ab41525, RRID:AB_732609 |
| Rabbit monoclonal anti-PKM1, 1:300 | Cell Signaling Technology | Cat# 7067, RRID: AB_2715534 |
| Rabbit monoclonal anti-PKM2, 1:300 | Cell Signaling Technology | Cat# 4053, RRID: AB_1904096 |
| Mouse monoclonal anti-Cytochrome C, 1:300 | Molecular Probes | Cat# A-6403, RRID: AB_221582 |
| Rabbit polyclonal anti-Cone arrestin, 1:300 | Sigma-Aldrich | Cat# AB15282; RRID:AB_1163387 |
| Rabbit polyclonal anti-Citrate synthase, 1:300 | GeneTex | Cat# GTX110624, RRID: AB_1950045 |
| Rabbit polyclonal anti-ATP synthase gamma, 1:300 | GeneTex | Cat# GTX114275S, RRID: AB_10726795 |
| Rabbit polyclonal anti- Fumarate hydratase, 1:300 | GeneTex | Cat# GTX109877, RRID: AB_1950283 |
| Rabbit polyclonal anti-Taurine, 1:300 | Abcam | Cat# ab9448, RRID: AB_307261 |
| Rabbit monoclonal anti-Alanine transaminase, 1:300 | Abcam | Cat# ab202083, RRID:AB_2915976 |
| Rabbit polyclonal anti-SUCLG1, 1:300 | Novus Biologicals | Cat# NBP1-32728, RRID: AB_2286802 |
| Mouse monoclonal anti-Glutamine synthetase, 1:1000 | Abcam | Cat# MAB302, RRID: AB_2110656 |
| Rabbit polyclonal anti-Glutaminase C (GAC), 1:300 | GeneTex | Cat# GTX131263, RRID: AB_2886452 |
| Rabbit polyclonal anti-PCK1, 1:300 | Affinity Biosciences | Cat# DF6770, RRID:AB_2838732 |
| Rabbit monoclonal anti-PCK2, 1:300 | Novus Biologicals | Cat# NBP2-75610  RRID: AB_2915974 |
| Peanut agglutinin (PNA), 1:1000 | Vector laboratories | Cat# FL-1071, RRID:AB_2315097 |
| Rabbit polyclonal anti-Aspartate aminotransferase-1, 1:300 | Abcam | Cat# ab221939,  RRID: AB_2915980 |
| Rabbit polyclonal anti-FABP-1 (Aspartate aminotransferase-2), 1:300 | Abcam | Cat# ab153924, RRID:AB_2915981 |

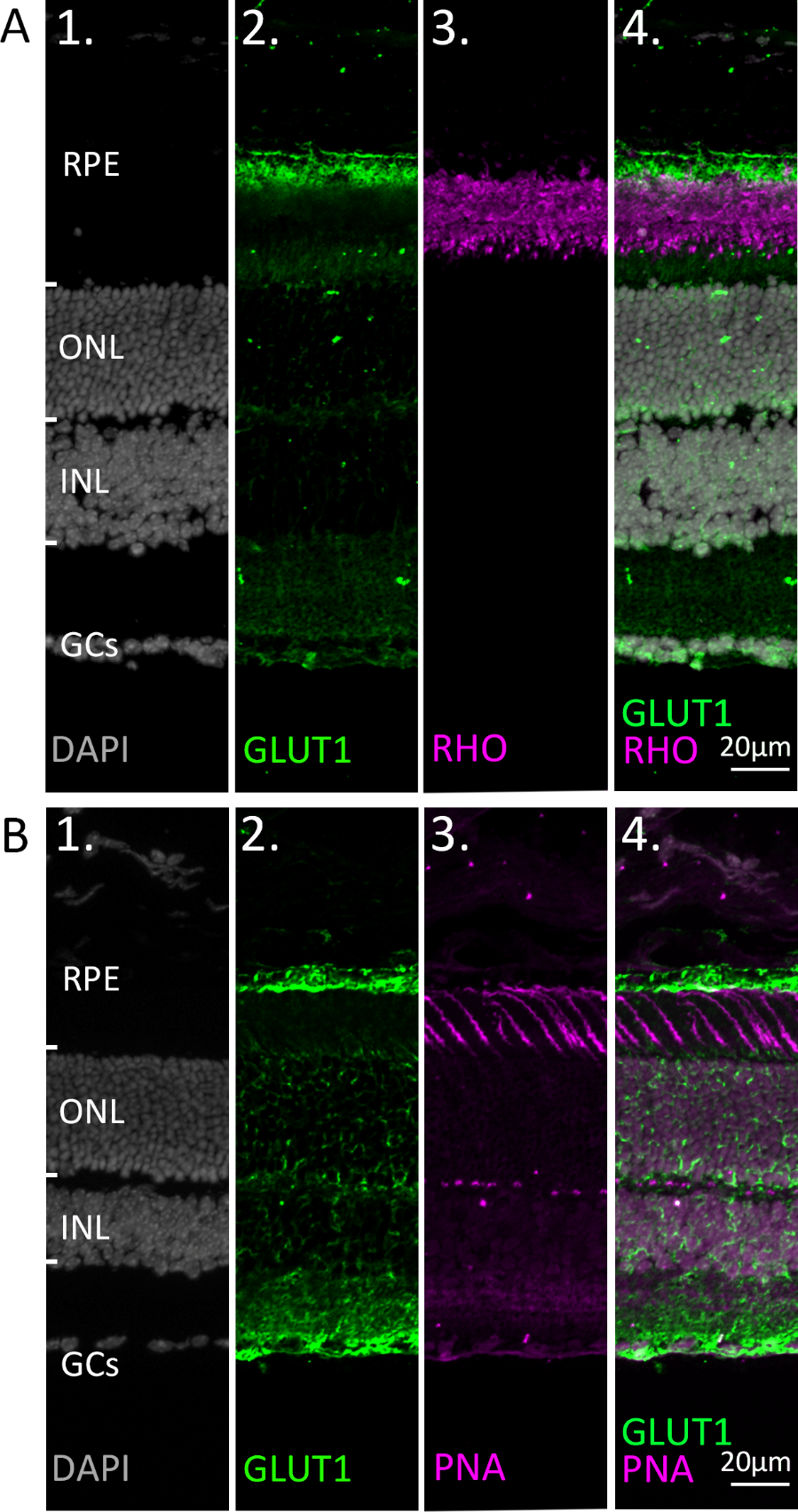

**Fig. S1. Co-staining of GLUT1 with markers for rod and cone photoreceptors.** Immunostaining for glucose transporter-1 (GLUT1; green). DAPI (grey) was used as nuclear counterstain. (A) Co-staining with the rod outer segment marker rhodopsin (RHO; magenta). (B) Co-staining with the cone marker peanut agglutinin (PNA; magenta). Note that while the retinal pigment epithelium (RPE) expresses high levels of GLUT1, both rod and cone photoreceptors appear to be devoid of GLUT1 expression. At high gamma values (B2) a staining suggestive of Müller cells is noticed. ONL = outer nuclear layer; INL = inner nuclear layer; GCs = ganglion cells.

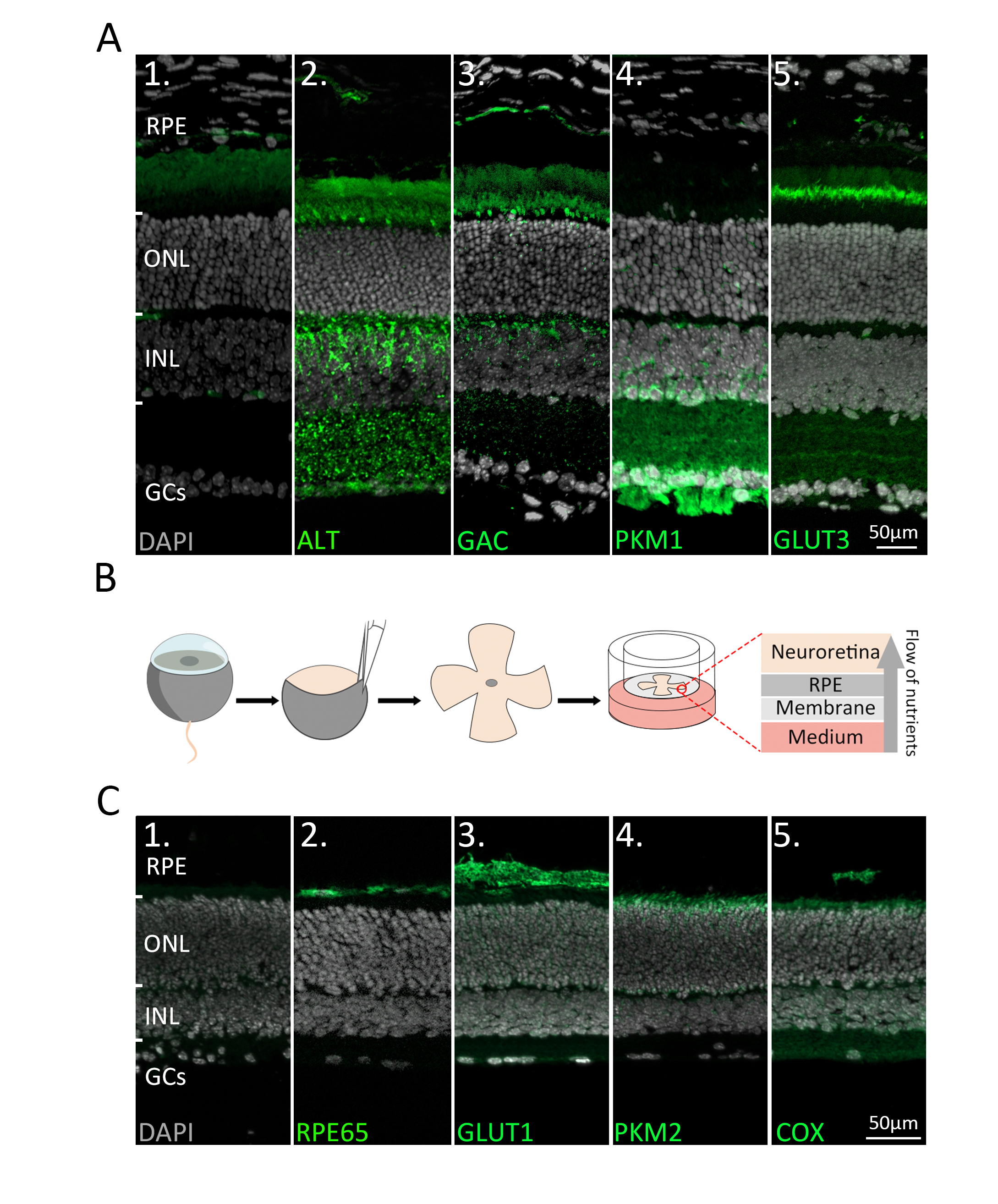

Fig. S2. Immunofluorescence staining of retinal enzymes and transporters *in vivo* and *in vitro*. (A) Protein expression (green) in the *in vivo* retina. DAPI (grey) was used as a nuclear counterstain. (1) Negative control. (2) Alanine transaminase (ALT) showed strong labelling on photoreceptor inner segments (IS) and inner nuclear layer (INL). (3) Glutaminase C (GAC) was expressed in IS and INL. (4) Pyruvate kinase M1 (PKM1) selectively labelled INL cells, synapses in the inner plexiform layer (IPL), and the ganglion cell layer (GCL). (5) Glucose transporter 3 (GLUT3) was strongly expressed on the basal side of IS and in discrete IPL sublamina. (B) Retinae were explanted at post-natal day (P) 9 and cultured until P15. During the culture period the retinal pigment epithelium (RPE) faced the culturing membrane so that nutrients from the underlying medium had to pass through the RPE. (C) Protein expression (green) in the *in vitro* retina. (1) Negative control. (2) RPE65. (3) Glucose transporter-1 (GLUT1). (4) Pyruvate kinase M2 (PKM2). (5) Mitochondrial cytochrome oxidase (COX). Note that the cultured retina is somewhat thinner than in a living animal, yet the *in vitro* enzyme expression patterns are comparable to the *in vivo* situation (*cf.* Figure 1A). ONL = outer nuclear layer; GCs = ganglion cells.

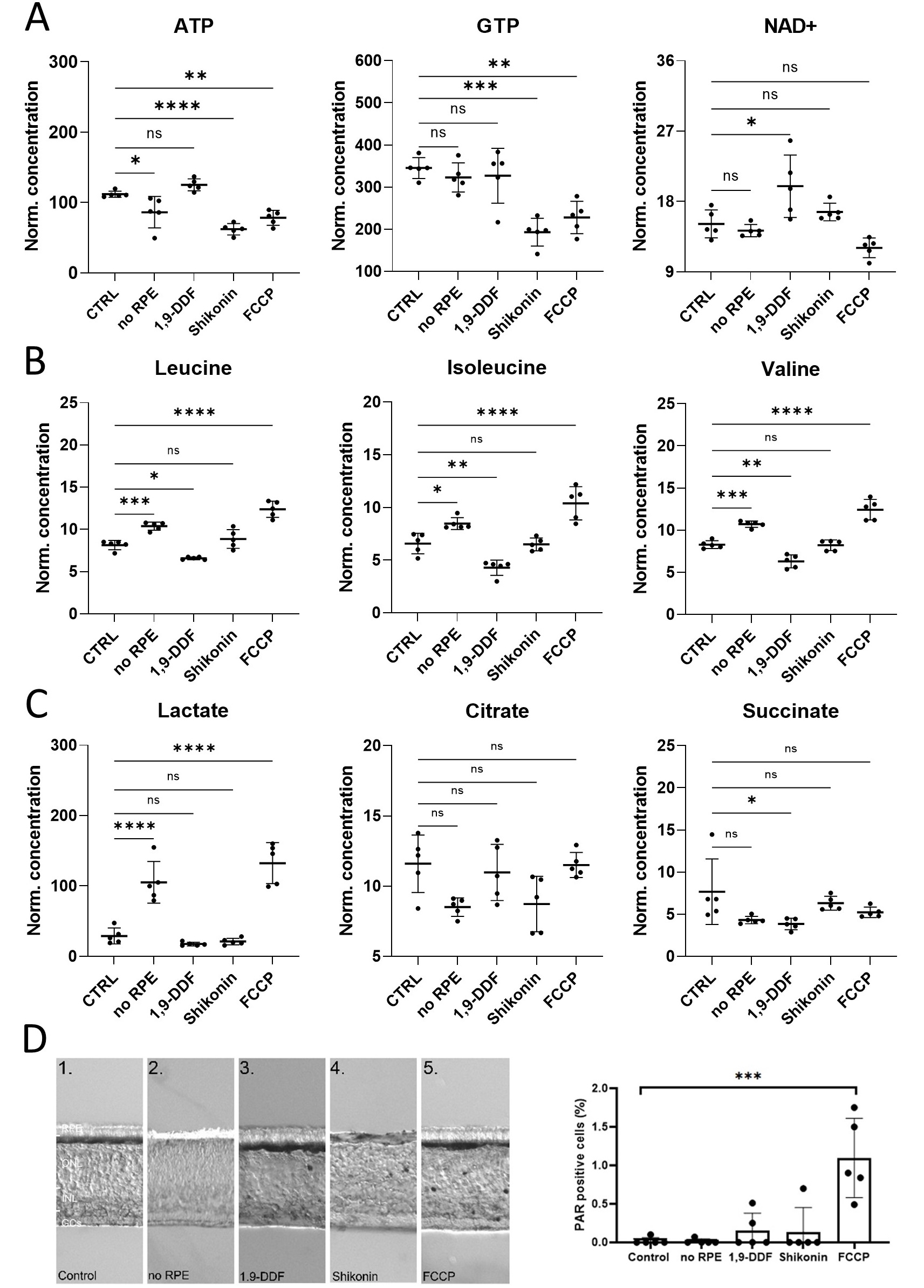

Fig. S3. Retina characterization by metabolomics and immunostaining. (A) Energy metabolism related metabolites: Overview box plots. (B) Box plots for branched chain amino acids (BCAA). (C) Krebs-cycle and glycolysis-associated metabolite box plots. Data represented as individual data points in µM concentration range with mean ± SD. (D) Immunodetection of poly-ADP-ribose (PAR) in the five experimental conditions. The bar graph shows the percentage of PAR positive cells in the outer nuclear layer (ONL). Statistical testing: One-way ANOVA with Tukey’s post-hoc test. For simplicity only *p*-values for comparison to control are shown: **** < 0.0001, *** < 0.001, ** < 0.01 * < 0.05.

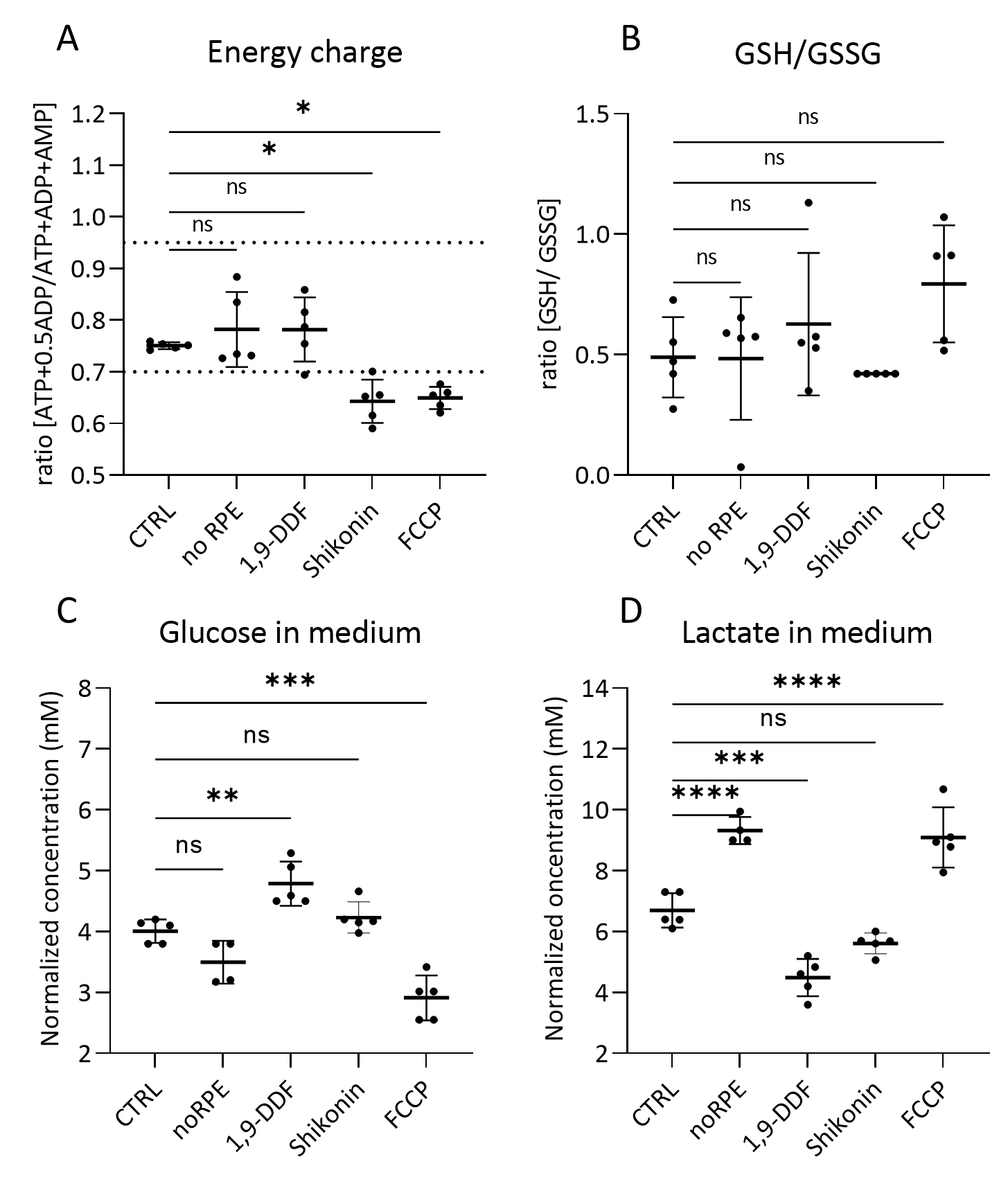

**Figure S4. Retinal tissue energy status, glutathione ratio, glucose, and lactate concentrations in medium.** Comparison of metabolism-related parameters across the five different experimental conditions. (A) The calculated ratio of the adenylate energy charge in the retina. The dotted lines indicate the energy charge window typical for most healthy cells (0.7 – 0.95). (B) Ratio of the glutathione (GSH)/glutathione disulfide (GSSG) in retina tissue. (C) Glucose concentration in the medium at the last day of retina explant culture (*i.e.* at P15). (D) Lactate concentration in P15 explant culture medium samples. Statistical testing: one-way ANOVA with Tukey’s post-hoc test. Data represented as individual data points as a metabolite concentration ratio (A, B) or in mM concentration range (C, D) with mean ± SD. Asterisks indicate significance levels; *p*-values: **** < 0.0001, *** < 0.001, ** < 0.01 * < 0.05.

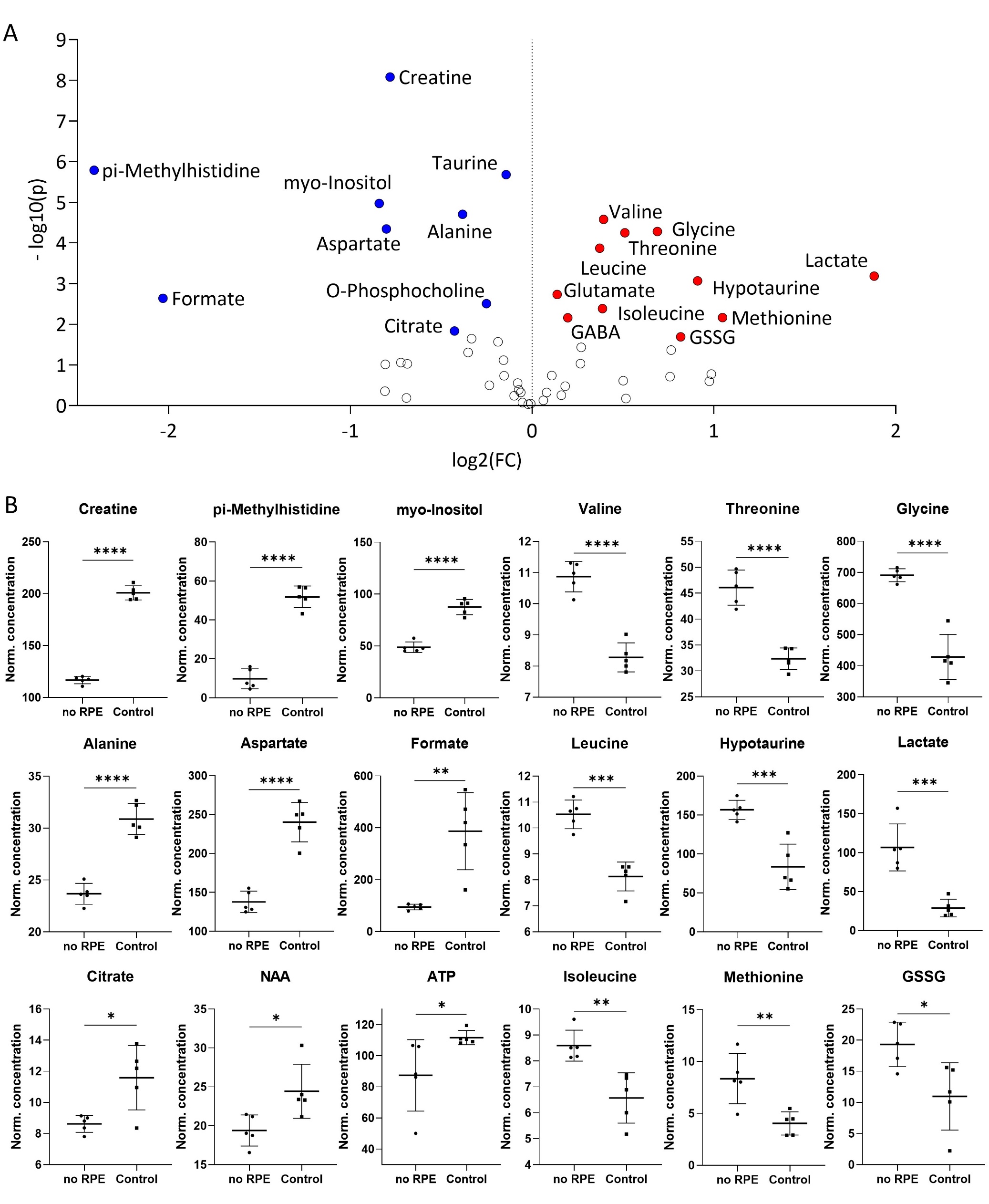

**Fig. S5.** **Metabolomic comparison of no RPE condition *vs*. control.** (A) Volcano plot illustrating the most significant metabolite changes. log_2_ (fold change; FC) = log_2_ (mean metabolite concentration in control) - log_2_ (mean metabolite concentration in no RPE group). (B) Individual metabolite box plots for FC > 1.2 and raw *p*-value < 0.05. Statistical testing: Student´s unpaired, two-tailed, *t*-test; *p*-values: **** < 0.0001, *** < 0.001, ** < 0.01 * < 0.05. Data represented as individual data points in µM concentration range with mean ± SD.

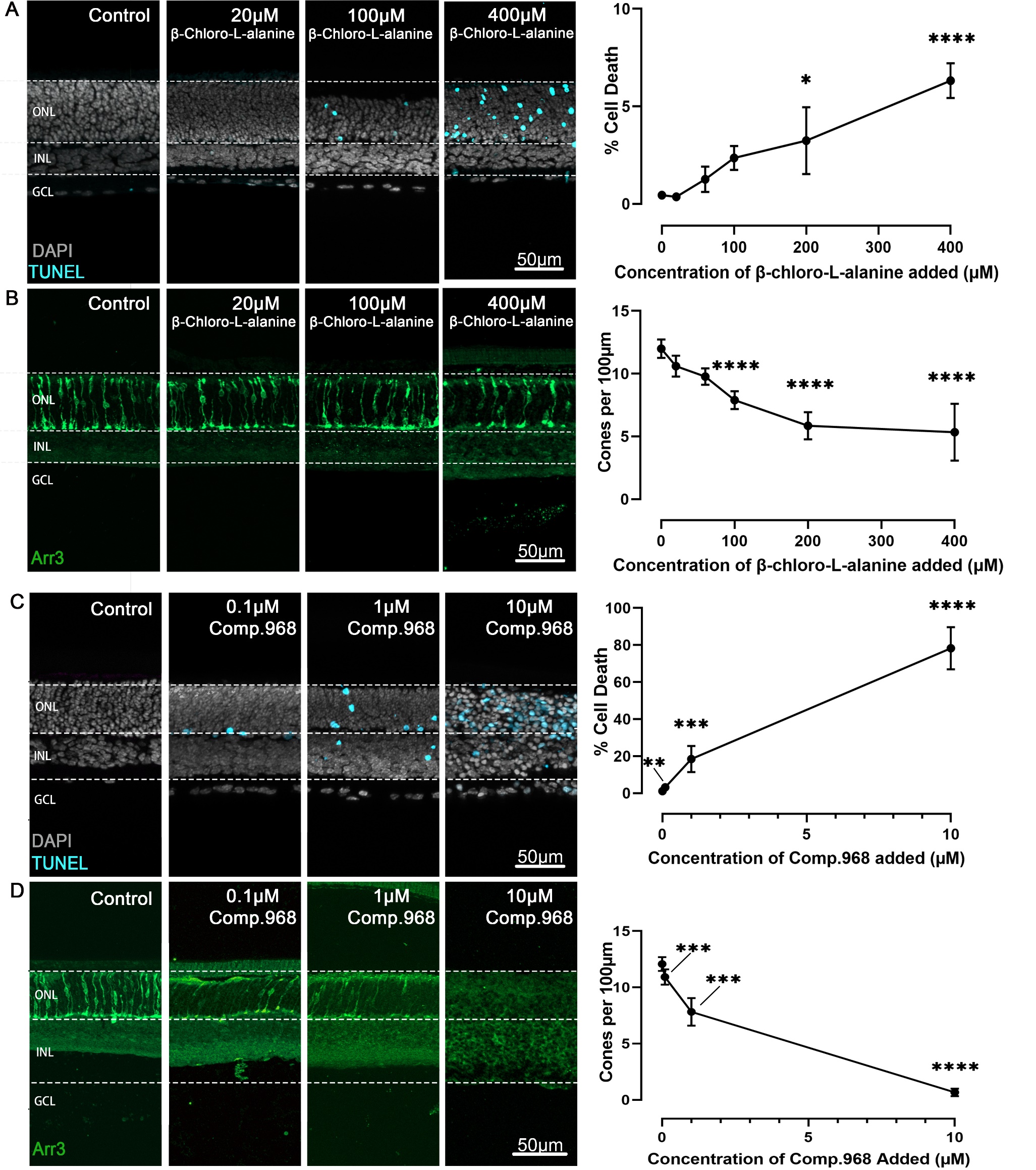

**Fig. S6.** **Inhibition of alanine transaminase and glutaminase C dose-dependently reduce photoreceptor viability.** Organotypic retina explants explanted at post-natal day (P) 9 were treated from P11 to P15 with inhibitors for either alanine transaminase (ALT) or glutaminase C (GAC). Left image panels display retinal cross-sections; graphs on right show quantifications of dying cells as evidenced by the TUNEL assay (cyan) or cone photoreceptors as indicated by cone arrestin-3 (Arr3) immunostaining (green); DAPI (grey) was used as nuclear counterstain. (A) ALT inhibition with ß-chloro-alanine selectively increased photoreceptor cell death in the outer nuclear layer (ONL) in a concentration-dependent manner, and (B) significantly decreased cone viability. (C) Inhibition of GAC with compound 968 dose-dependently increased photoreceptor cell death, and (D) significantly reduced cone viability. Note that both ALT and GAC inhibition left the inner retina essentially intact. INL = inner nuclear layer; GCL = ganglion cell layer; scale bare = 50 µm. Statistical comparison using one-way ANOVA with Tukey’s multiple comparisons test; *p*-values: **** < 0.0001, *** < 0.001, ** < 0.01, * < 0.05.

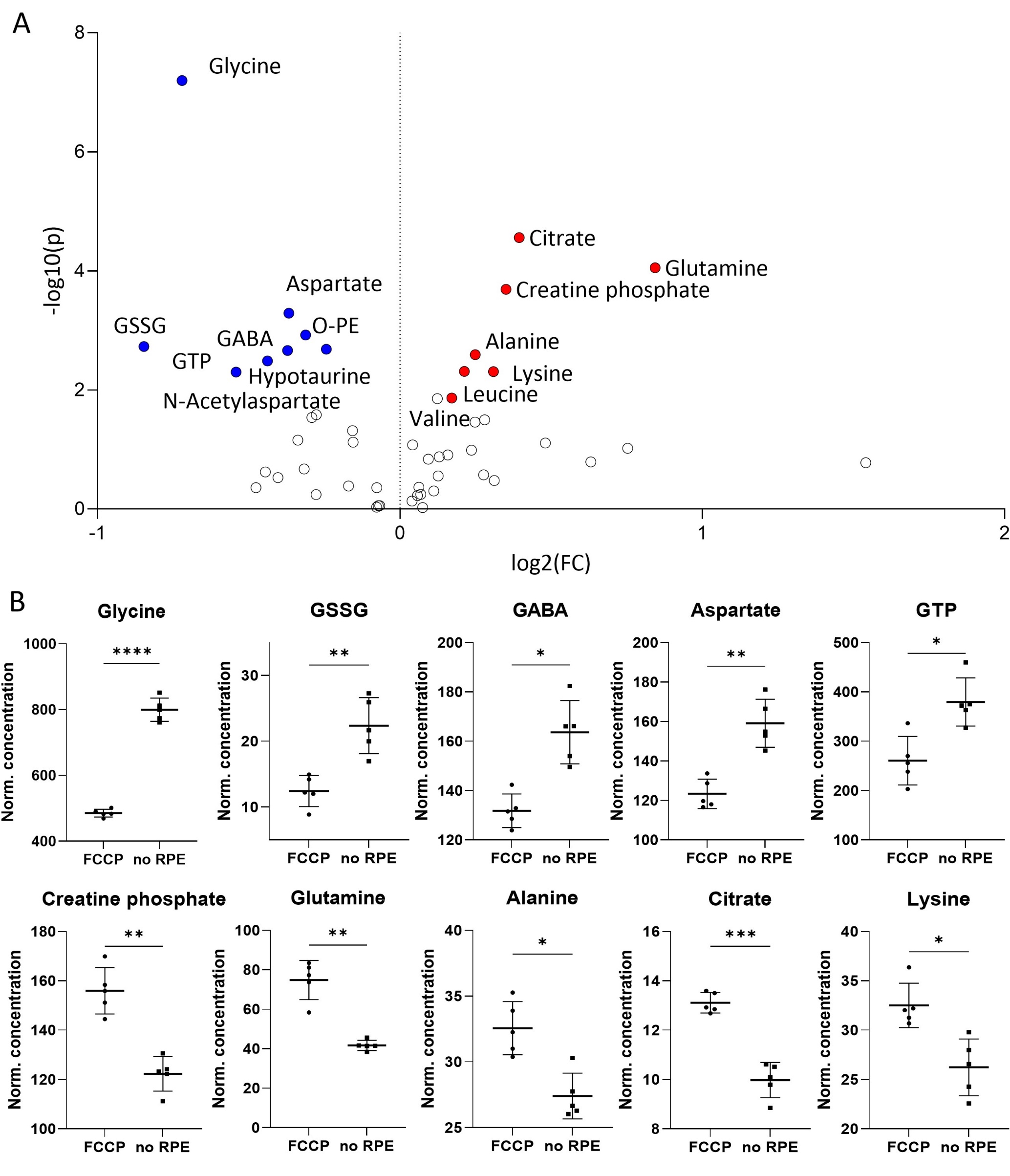

**Fig. S7. Metabolomic comparison of** **FCCP *vs*. no RPE.** (A) Volcano plot illustrating the most significant metabolic changes. log_2_ (fold change; FC) = log_2_ (mean metabolite concentration in FCCP group) - log_2_ (mean metabolite concentration in no RPE group). (B) Individual metabolite box plots for FC > 1.2 and raw p value < 0.05. Statistical testing: Student´s unpaired, two-tailed, *t*-test; *p*-values: **** < 0.0001, *** < 0.001, ** < 0.01 * < 0.05. Data represented as individual data points in µM concentration range with mean ± SD.

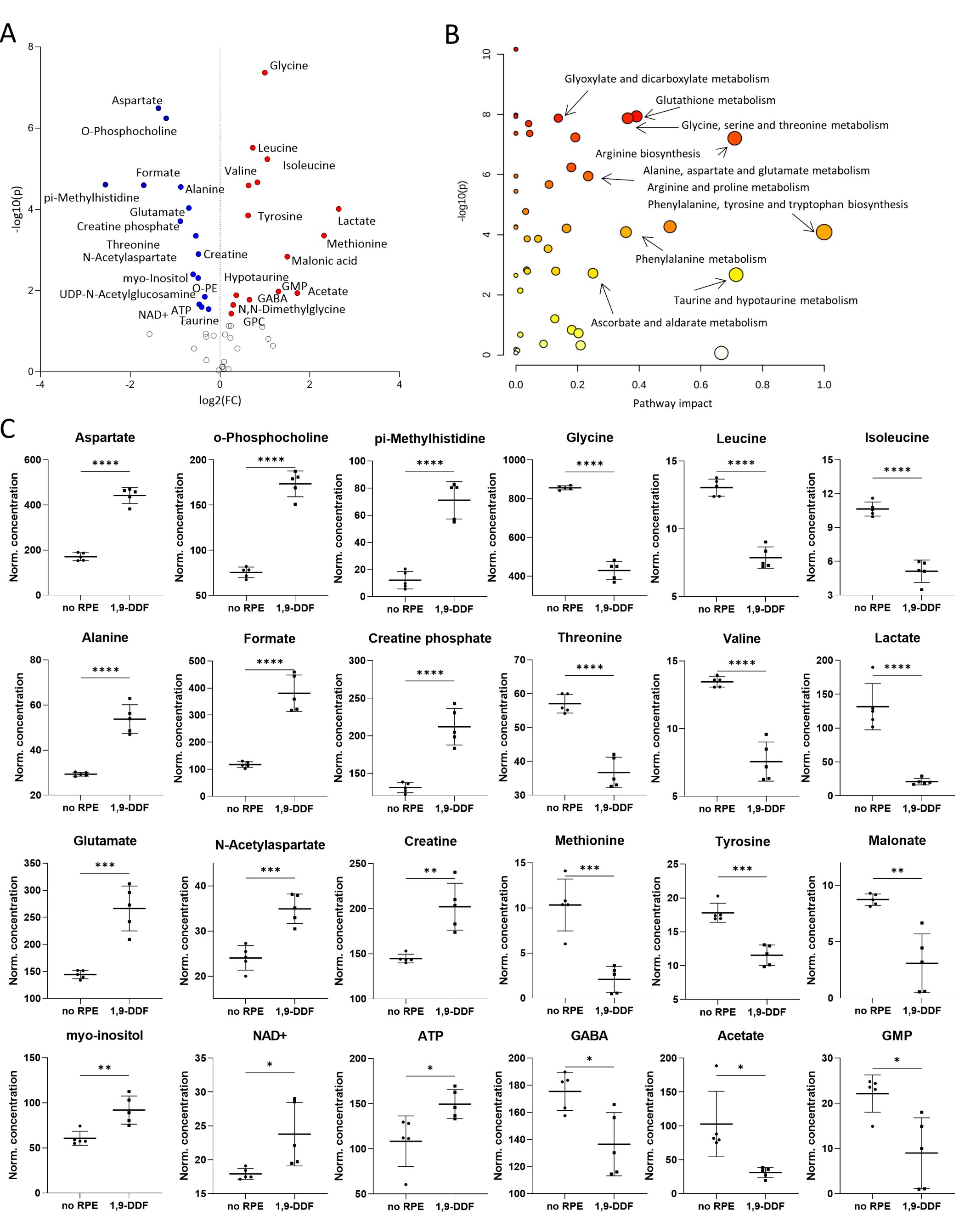

**Fig. S8.** **Metabolomic comparison of** **no RPE *vs*. 1,9-DDF.** (A) Volcano plot illustrating the most significant metabolic changes. log_2_ (fold change; FC) = log_2_ (mean metabolite concentration in no RPE group) - log_2_ (mean metabolite concentration in 1.9-DDF group). (B) KEGG database-based pathway analysis with the most affected metabolic pathways. (C) Individual metabolite box plots for FC > 1.2 and raw p value < 0.05. Statistical testing: Student´s unpaired, two-tailed, *t*-test; *p*-values: **** < 0.0001, *** < 0.001, ** < 0.01 * < 0.05. Data represented as individual data points in µM concentration range with mean ± SD.

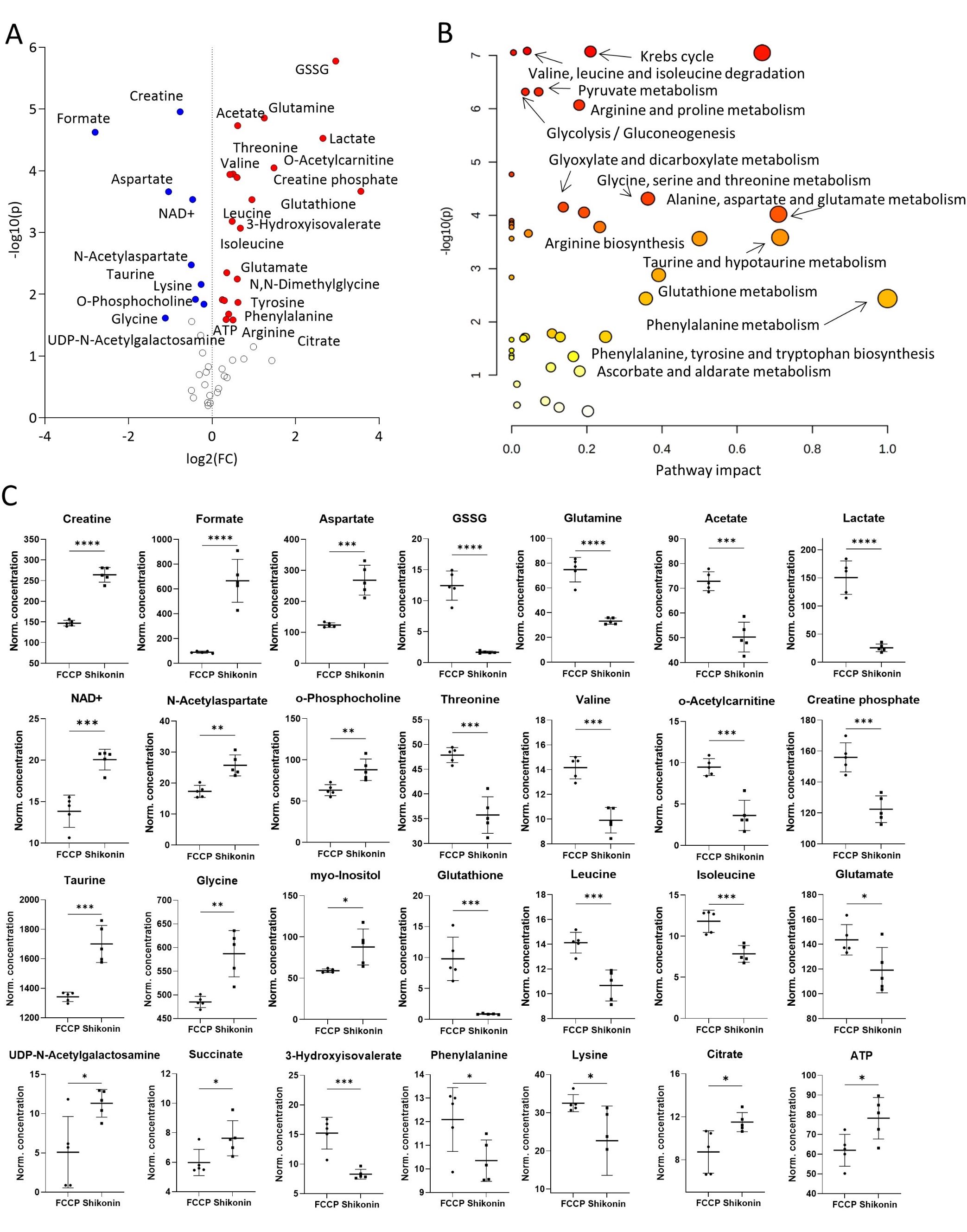

**Fig. S9.** **Metabolomic comparison of** **FCCP *vs*. Shikonin.** (A) Volcano plot with the most significant metabolic changes illustration. log_2_ (fold change; FC) = log_2_ (mean metabolite concentration in FCCP group) - log_2_ (mean metabolite concentration in Shikonin group). (B) KEGG database-based pathway analysis with the most affected metabolic pathways. (C) Individual metabolite box plots for FC > 1.2 and raw *p*-value < 0.05. Statistical testing: Student´s unpaired, two-tailed, *t*-test; *p*-values: **** < 0.0001, *** < 0.001, ** < 0.01 * < 0.05. Data represented as individual data points in µM concentration range with mean ± SD.

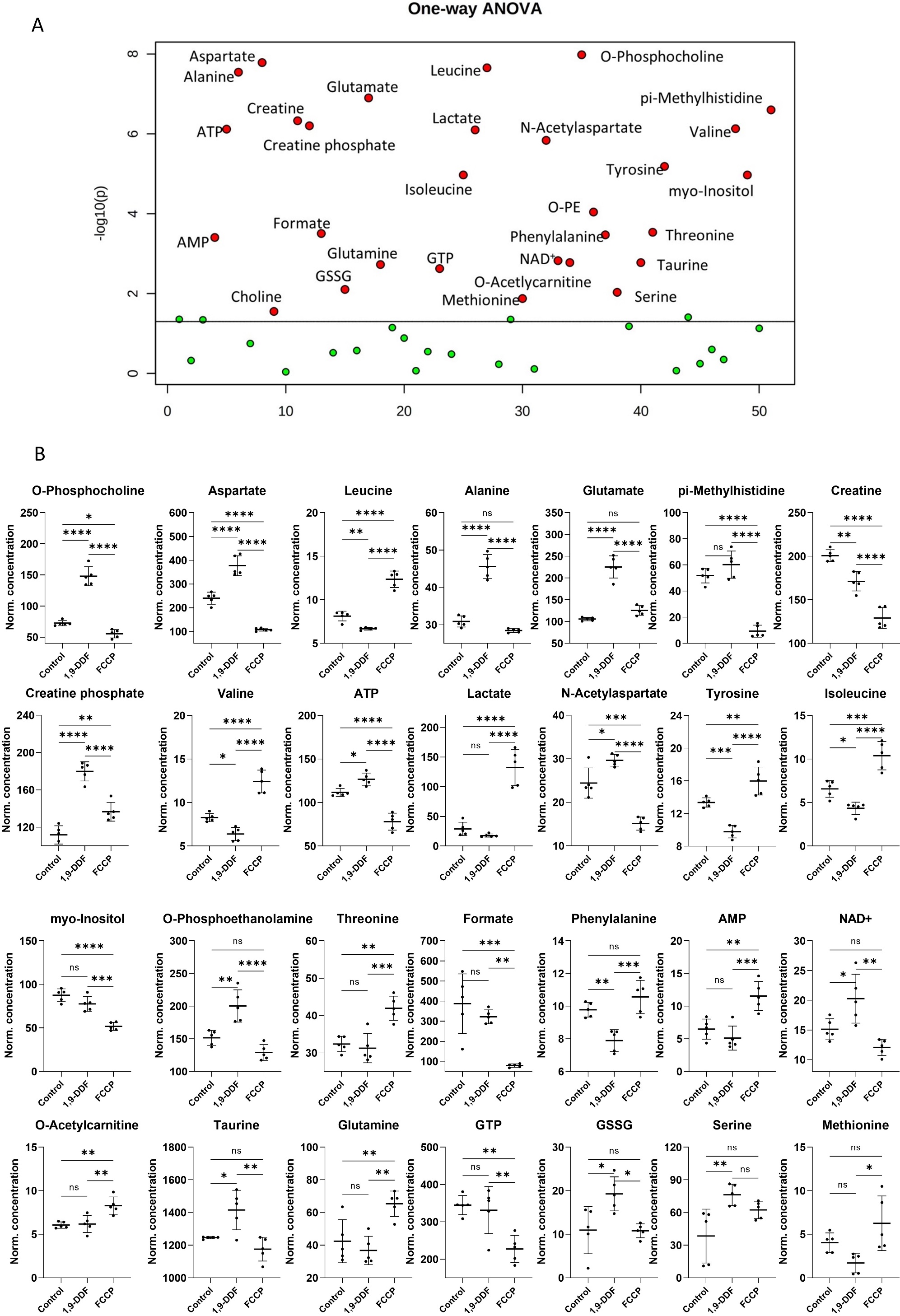

**Fig. S10.** **Metabolomics analysis of control *vs*. 1,9-DDF vs. FCCP three-way comparison.** (A) One-way ANOVA result visualization plot displaying the most significant metabolite concentration changes based on the 3-group analysis. The x-axis labels a numerical list of quantified and analyzed metabolites in alphabetical order. (B) Individual metabolite box plots. Statistical analysis: one-way ANOVA, Fisher’s LSD post-hoc analysis, *p*-values: **** < 0.0001, *** < 0.001, ** < 0.01 * < 0.05. Data shown as individual data points in µM concentration range with mean ± SD.
